# Msp300 Links Autophagy to Nuclear Morphodynamics

**DOI:** 10.64898/2026.11.02.763579

**Authors:** Shristi Sharma, Ioannis P. Nezis

## Abstract

Autophagy is a fundamental cellular process that maintains homeostasis and enables adaptation to stress by degrading damaged or superfluous components. While its core machinery is well characterized, the interplay between autophagy and nuclear architecture remains poorly understood. In this study, we identify the LINC complex protein Msp300 as a novel interactor of the autophagy protein Atg8a in *Drosophila melanogaster*. Msp300 belongs to the KASH (Klarsicht/ANC-1/Syne Homology) domain protein family, known for its role in nuclear positioning through interactions with the cytoskeleton. The KASH domain anchors Msp300 in the outer nuclear membrane, where it pairs with SUN domain proteins across the nuclear envelope to form the LINC complex, a mechanical bridge between the nucleoskeleton and cytoskeleton. Yeast two-hybrid screening revealed an interaction between Atg8a and Msp300 isoforms E and G, mediated by a canonical LC3-interacting region (LIR). Localization studies show Msp300 associates with Atg8a within the cytoplasm and at the nuclear periphery under both basal and elevated autophagy conditions. Functional perturbations through stress assays indicate that this interaction influences event in and around nucleus, establishing a link between autophagy and nuclear architecture. By exploring the role of Msp300 isoforms E and G in shaping not only nuclear organization but potentially endoplasmic reticulum, through autophagy machinery, this study introduces a new dimension to nuclear envelope biology and opens opportunities to explore how steady state homeostasis, stress adaptation and cytoskeletal connectivity are coordinated at the nuclear periphery.

## Introduction

Autophagy, derived from the Greek words *auto* (self) and *phagy* (to eat), was first described by Christian de Duve in 1963 (De Duve and Wattiaux, 1966). It is a conserved cellular degradation process that eliminates damaged or superfluous cellular components including proteins, lipids, nucleic acids, and organelles-via lysosome-mediated breakdown. This process is essential for maintaining cellular homeostasis, supporting development and differentiation, and enabling survival under stress conditions (Aman *et al*., 2021). Physiological and pathological implications of autophagy extend way beyond these roles (Morishita and Mizushima, 2019). Among the various forms of autophagy, macroautophagy is the most prominent. It involves the formation of double-membrane structures called autophagosomes, which engulf cytoplasmic cargo and subsequently fuse with lysosomes for degradation (Mercer *et al*., 2018). This process is tightly regulated by a suite of autophagy-related genes (Atgs), Atg8a, Atg12, and Ref (2) P to name a few. Notably, *Atg8a*, a ubiquitin-like protein, undergoes lipidation with phosphatidylethanolamine (PE), allowing it to associate with autophagic membranes and facilitate phagophore expansion and closure (Jipa *et al*., 2021). Autophagy is not merely a bulk degradation pathway instead it can be highly selective. This selectivity is mediated by selective autophagy receptors (SARs), which bind specific cargo and interact with Atg8-family proteins via conserved LC3-interacting region (LIR) motifs (Johansen and Lamark, 2011). Lipdated-Atg8 appears to be a central factor in mediating cargo selectivity through direct interactions with SARs (Farré and Subramani, 2016). The cargo once recognized is attached to the phagophore and ultimately degraded by fusion with lysosome.

To further elucidate the molecular landscape of autophagy, we worked to identify novel proteins interactors of key autophagy Atg8-family proteins in *Drosophila*. A yeast-two-hybrid screening was performed by hybrigenics services (https://www.hybrigenics-services.com) in 3rd instar larvae and head of *Drosophila* which uncovered a novel Atg8a interactor- a nuclear envelope LINC complex protein, Msp300. Previous studies have shown that Msp300 contributes to myonuclear spacing, organelle distribution, and cytoskeletal organization (Elhanany-Tamir *et al*., 2012). More recently, Msp300 has been implicated in autophagy, with its loss leading to accumulation of un-degraded aggregates marked by autophagy components such as Ref (2)P and Atg8a, suggesting a block in autophagic flux (Elhanany-Tamir *et al*., 2012). This study also registers that Msp300 localizes throughout the ER network and proposes that Msp300 acts as a structural and functional adapter between the cytoskeleton and organelles, influencing both autophagy and nuclear positioning (*et al*., 2025). Despite these findings, the mechanistic basis of Msp300’s role in autophagy, particularly its interaction with Atg8a, remains uncharacterised and poorly understood.

In this study, we hypothesize that that Msp300 interacts with Atg8a via a LIR motif contributing to autophagosome maturation and nuclear and ER architecture regulation under basal conditions and in stress. Like all other cellular organelles, the nucleus and ER undergoes constant changes, and dynamic reorganisation may it be for cellular transport or for the quality control. Nucleus houses the master regulator of the cell which is also prone to stress and damage. Nucleophagy is a form of autophagy specific to the nucleus and its components, recycling damaged nuclear components and managing cellular stress. In *Drosophila melanogaster*, autophagic turnover of the nucleus and nuclear components has been reported, but no receptor analogous to yeast Atg39 (Mochida *et al*., 2015) or mammalian UBC9 (Lee *et al*., 2020; Allegretti *et al*., 2020) has been reported to date. Similarly in case of ER, till date very less is known about regulation and receptors of ER-phagy. Rtnl1 (Reticulon-like 1), Keap1(Kelch-like ECH-associated protein 1), and PINK1(PTEN-induced kinase 1) have been recently identified as ER-phagy receptors for developmental autophagy (Wang *et al*., 2023). Building on the identified receptors partnership, Vps13(Vacuolar Protein Sorting 13) was additionally found as a vital genetic partner along with Atl and Rtnl1 under starvation autophagy. Here, we found a novel interaction between Atg8a and Msp300 Isoform E/G. We showed the role of Msp300 Isoform E/G in maintaining autophagic flux and explored the potential of Msp300 to be an autophagy receptor for ONM and ER, with a functional LIR motif. This study lays the foundation for future work with an Msp300-mcherry-gfp tandem reporter that could establish Msp300 as a micro-ONM- phagy receptor and will be a stepping stone towards exploring its role in ER-phagy.

## Methodology

### Yeast- two-hybrid Interaction Study

Yeast two-hybrid (Y2H) interaction studies to investigate the interaction between Msp300 and Atg8a were performed by Hybrigenics Services (Evry, France) using the ULTImate Y2H™ screening platform. *Drosophila melanogaster* full-length Atg8a (aa 1-121) was used as the bait protein. Atg8a was cloned into the pB27 bait vector to generate an N-LexA-Atg8a-C fusion (bait construct hgx4666v1). The bait was screened against *D. melanogaster* cDNA libraries derived from third-instar larvae and adult fly heads. pB7 was the Gal4 activation domain vector in which Msp300 E target (hgx6778v1, hgx6779v1 and hgx6780v1) and Msp300 G target (hgx6976v1) were cloned into pP14 Gal4 activation domain vector for one-on-one interaction. In case of Msp300 G screening, methionine is used to regulate expression of the prey fusion. Under the MET25 promoter and in medium lacking methionine (DO3-MET) prey fragment’s expression is induced, while medium lacking tryptophan/leucine (DO2) serves as a growth/plasmid-presence control. The screens were performed using Hybrigenics cell to cell mating-based Y2H procedure, in which interaction between the Atg8a bait and a prey protein activates the HIS3 reporter.

Colonies were selected by growth under tryptophan, leucine and histidine-deficient conditions (DO3) for Msp300 E and in histidine-deficient conditions along with Methionine-deficient conditions for Msp300 G.expression of the prey fusion. Under the MET25 promoter and in medium lacking methionine (DO3-MET) prey fragment’s expression is induced, while medium lacking tryptophan/leucine (DO2) serves as a growth/plasmid-presence control. The screens were performed using Hybrigenics cell to cell mating-based Y2H procedure, in which interaction between the Atg8a bait and a prey protein activates the HIS3 reporter. Colonies were selected by growth under tryptophan, leucine and histidine-deficient conditions (DO3) for Msp300 E and in histidine-deficient conditions along with Methionine-deficient conditions for Msp300 G.

### AlphaFold 3 protein-protein interaction prediction

AlphaFold 3 was used to investigate the potential interaction between Atg8a and Msp300 isoform E and G. - Msp 300 is too large a protein to model as a full-length complex, and AlphaFold-Multimer-type predictions lose sensitivity with long sequences), particularly for interactions mediated by short linear motifs in disordered regions (Lee *et al*., 2024). We therefore divided Msp300 E into seven overlapping fragments and Msp300 G into ten overlapping fragments of about 1,500 residues (150-residue overlap between corresponding fragment) and co-folded each with Atg8a (121 aa residues) using AlphaFold Server (AlphaFold 3; Abramson *et al*., 2024). The result “.zip” file was expanded and interpretated using AlphaBridge.

Initially, all the Msp300 isoform E and G target fragments of approximately 1,500 amino acids were modelled with Atg8a to identify a possible interaction site. Once the interaction was identified, a shorter fragment of 140 amino acids encompassing the predicted interaction region was selected for further analysis the target was contained in intrinsically disordered regions of the protein. To investigate the specificity of the predicted interaction, a scrambled version of the fragment and another protein Ultrabithorax isoform F, with highly disordered region was also analysed as a computational control. Additionally, selected amino acids previously predicted by Y2H analysis were mutated in same fashion as LIR mutant of Y2H analysis, and the resulting sequence was modelled to assess whether the predicted interface was retained or altered.

Fragment trimming and motif scrambling were performed in Google Colaboratory (https://colab.research.google.com). The motif was scrambled by random shuffling (seeds 1 and 2), keeping amino-acid composition and flanking sequence unchanged and requiring at most 30% positional identity to the original while the sequence scrambled had 9% similarity to new scrambled motif. AlphaBridge helped to identify predicted protein-protein interfaces and assess the associated interface scores, including ipTM, piCS and the AlphaBridge score.

### Fly husbandry and generation of Transgenic flies

Flies used in experiments were reared at 25°C, 70% relative humidity, and raised on a cornmeal-based diet containing 42g Inactive Yeast (Dutscher Ltd, 789126), 60g Cornmeal (Quaker®, 00030000570203), 130g Sucrose, 6g Agar (Thermo Fisher Scientific Inc., 214010), 10% w/v Nipagin, per litre of water. All genotypes of flies sourced from Drosophila sourcing centres-BDSC and VDRC, or genotypes acquired as kind gift from other researchers are listed **(Table 1).**

**Table 1.** Fly Stocks used in this study.

| **Name** | **Genotype** | **Source** |
|---|---|---|
| Isogenic control used to create Msp300 E/G LIR mutant | W[1118] | WellGenetics |
| Msp300<sup>complete mutant</sup> | W[1118]; Msp300complete neoFRT40/Cyo, GFP | Gifted from Timothy Megraw, University of Florida |
| Msp300 GFP (BL 59757) | y[1] w[67c23]; Mi{PT-GFSTF.0}Msp300[MI00111-GFSTF.0]/ CyO | Bloomington Stock Center |
| Msp300 E/G LIR mutant | W[1118]; Msp300-RE Y8968A I8971A CRISPR/ CyO | Created by WellGenetics |
| Msp300 E/G Null mutant | W[1118]; Msp300-RE Y8968A I8971A CRISPR{PBacDsRed} / CyO | Created by WellGenetics |
| Atg8a RNAi Mutant (109654) | hsFLP; UAS Dcr2; act CDZ Gal4, UAS-GFP Nls/UAS Atg8a RNAi | Vienna *Drosophila* Resource Center |
| Atg8<sup>Tro- Gal4</sup>(Atg8a mutant) | FRTAGA, ATG8a<sup>Tro-GAL4</sup>/ FM7, GFP; +/+; +/+ | Gifted from Gábor Juhász, University of Budapest. |
| Atg8<sup>KG</sup> (Atg8a mutant) | y P(hsFLP) Atg8a [KG07569] | Gifted from Gábor Juhász, University of Budapest. |
| FRT19A Flippase Stock | hsFLP, FRT 19A, ubiGFP; +/+; +/+ | Gifted from Gábor Juhász, University of Budapest. |
| FRT40A Flippase Stock | YW; ubi-mRFP.Nls neoFRT40A / Sm6a | Gifted from Timothy Megraw, University of Florida |
| Heat Shock Flippase | hsFLP; UAS Dcr2; act CDZ Gal4, UAS-GFP Nls | Gifted from Péter Lőrincz Eotvos Lorand University |
| Tandem Atg8a Reporter | W[1118];Cg-Gal4/UAS-GFP m-Cherry-Atg8a | Created in-house |
| Flippout m-cherry Atg8a | hs-FLP/+; cg>CD2>GAL4, UAS-mCherry-Atg8a/+ | Created in-house |
| CgGal4 m-cherry Atg8a | W[1118];Cg-Gal4/UAS-m-Cherry-Atg8a | Created in-house |

**Table 2.**
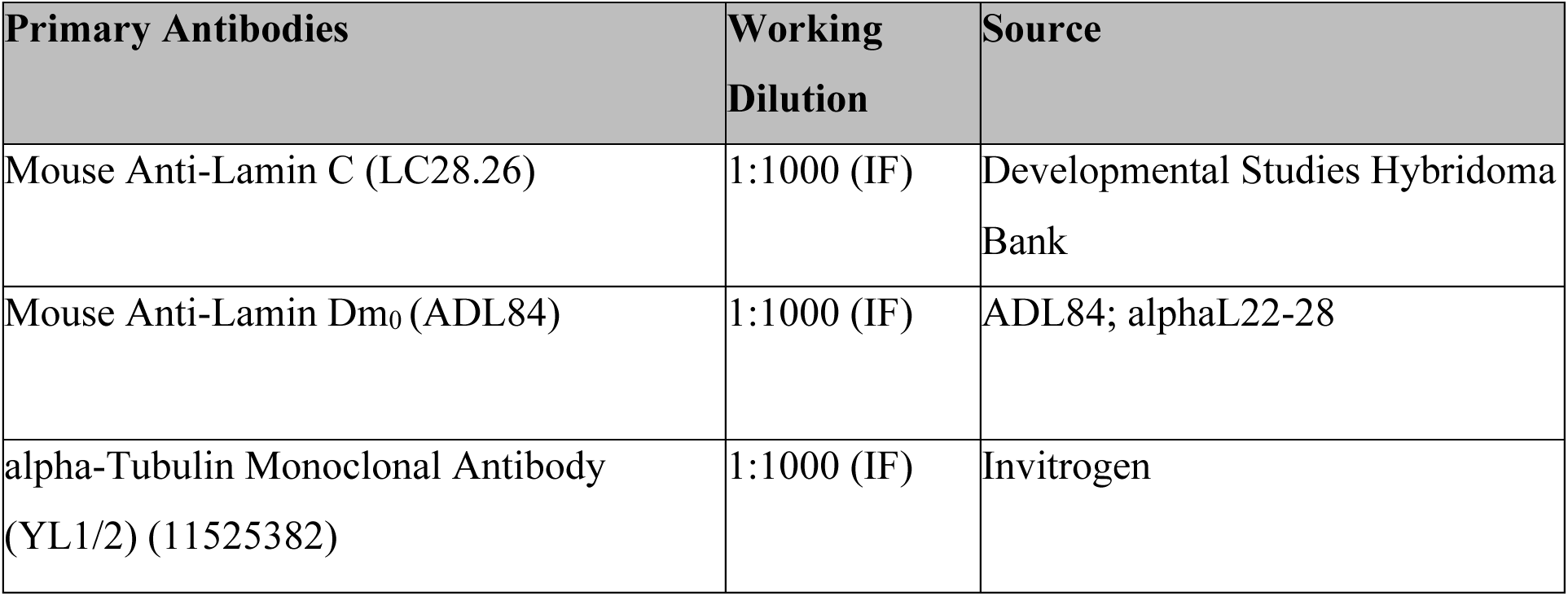
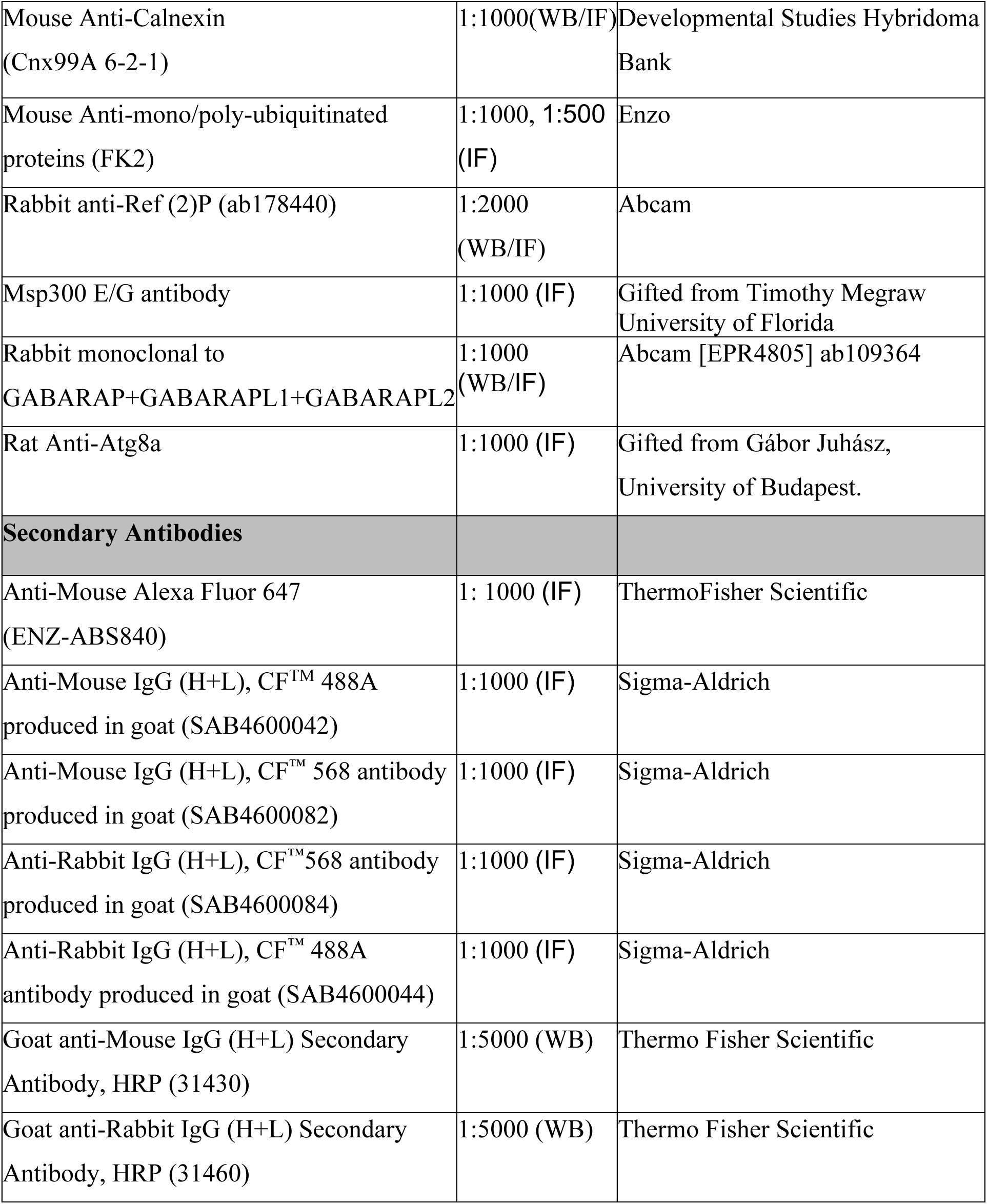
Antibodies and Dyes used in this study for immunohistochemistry and staining experiments.

| Primary Antibodies | Working Dilution | Source |
|---|---|---|
| Mouse Anti-Lamin C (LC28.26) | 1:1000 (IF) | Developmental Studies Hybridoma Bank |
| Mouse Anti-Lamin Dm<sub>0</sub> (ADL84) | 1:1000 (IF) | ADL84; alphaL22-28 |
| alpha-Tubulin Monoclonal Antibody (YL1/2) (11525382) | 1:1000 (IF) | Invitrogen |
| Mouse Anti-Calnexin (Cnx99A 6-2-1) | 1:1000(WB/IF) | Developmental Studies Hybridoma Bank |
| Mouse Anti-mono/poly-ubiquitinated proteins (FK2) | 1:1000, 1:500 (IF) | Enzo |
| Rabbit anti-Ref (2)P (ab178440) | 1:2000 (WB/IF) | Abcam |
| Msp300 E/G antibody | 1:1000 (IF) | Gifted from Timothy Megraw<br>University of Florida |
| Rabbit monoclonal to GABARAP+GABARAPL1+GABARAPL2 | 1:1000 (WB/IF) | Abcam [EPR4805] ab109364 |
| Rat Anti-Atg8a | 1:1000 (IF) | Gifted from Gábor Juhász, University of Budapest. |
| **Secondary Antibodies** |  |  |
| Anti-Mouse Alexa Fluor 647 (ENZ-ABS840) | 1: 1000 (IF) | ThermoFisher Scientific |
| Anti-Mouse IgG (H+L), CF<sup>TM</sup> 488A produced in goat (SAB4600042) | 1:1000 (IF) | Sigma-Aldrich |
| Anti-Mouse IgG (H+L), CF™ 568 antibody produced in goat (SAB4600082) | 1:1000 (IF) | Sigma-Aldrich |
| Anti-Rabbit IgG (H+L), CF™568 antibody produced in goat (SAB4600084) | 1:1000 (IF) | Sigma-Aldrich |
| Anti-Rabbit IgG (H+L), CF™ 488A antibody produced in goat (SAB4600044) | 1:1000 (IF) | Sigma-Aldrich |
| Goat anti-Mouse IgG (H+L) Secondary Antibody, HRP (31430) | 1:5000 (WB) | Thermo Fisher Scientific |
| Goat anti-Rabbit IgG (H+L) Secondary Antibody, HRP (31460) | 1:5000 (WB) | Thermo Fisher Scientific |

Msp300 E/G LIR ^Y8968A I8971A^ CRISPR flies were created using CRISPR/Cas9-mediated genome editing by homology-dependent repair (HDR) using a guide RNA and a dsDNA plasmid donor, conducted by Wellgenetics Inc. (Taipei, Taiwan, China). gRNA1 sequence 5’-CTTCGATTGATTTCAGGTATCGCTT [CCG] and gRNA2 5’-CTTCGAAGATTAAGAGCAAGGATTT [CCG] were cloned in XP3-hsp70 Promoter. The PBacDsRed selection marker cassette comprises PiggyBac 3′ and 5′ terminal repeats, the artificial 3xP3 promoter (containing three tandem Pax-6 homodimer binding sites and an hsp70 TATA-like sequence), DsRed2, and the SV40 3′ UTR. This cassette was cloned into the pUC57-Kan vector to serve as the donor template for homology-directed repair. For genome editing, Msp300/CG42768-specific gRNAs and hs-Cas9 were supplied in DNA plasmids, along with the donor plasmid, which was microinjected into embryos of the W[1118] control strain. F1 flies carrying the selection marker 3xP3-DsRed were then validated through genomic PCR and sequencing. CRISPR introduced a double stranded break in the Msp300/ CG42768 gene, which was subsequently replaced by the PBacDsRed cassette. These Msp300 flies with PBacDsRed cassette were subsequently used in this study and referred to as Msp300 E/G Null flies because these flies have lost function copies of both the isoforms E and G. Finally, the PBacDsRed cassette was excised, effectively reconstituting the functional gene, which is now the Msp300 E/G LIR ^Y8968A I8971A^ mutant, expressed across the entire genome of the selected fly strains. Mutant flies were validated by genomic PCR and sequencing.

### Immunohistochemistry, Image Analysis and Statistics

Second-instar larvae were collected from fly-rearing tubes, washed thoroughly with water, and maintained in phosphate buffer saline (PBS; Merck & co., Inc., P4417) prior to dissection.

Tissues were dissected in ice-cold PBS under a stereomicroscope and fixed in 4% paraformaldehyde (Thermo Fisher Scientific Inc., 28908) for 30 min. Following fixation, tissues were permeabilized for 30 min in PBS containing 0.2% (v/v) Triton X-100 (Merck & co., Inc., T9284) and blocked for 1 h in blocking buffer-PBS containing 0.1% (w/v) bovine serum albumin (BSA) and 0.2% (v/v) Triton X-100. Tissues were incubated overnight at 4°C with primary antibodies **(Table. 2)** diluted in blocking buffer. Following three 10-min washes in PBS containing 0.1% Tween-20 (Thermo Fisher Scientific Inc., 85115), tissues were incubated with the appropriate secondary antibodies **(Table. 2)** for 2 h at room temperature. Samples were subsequently washed three times for 10 min in PBS containing 0.1% Tween-20 and once for 10 min in PBS.

For phalloidin (Thermo Fisher Scientific Inc., T7471(1:100) and T123799(1:1000)) or Nile red (1:1000; Thermo Fisher Scientific Inc., N1142) staining, tissues were incubated with the required dye following antibody staining for 30 minutes and washed once with PBS before mounting in vectashield mounting medium containing DAPI (Vector Laboratories, Inc., H-1200-10). For experiments involving dye staining alone, antibody incubation steps were omitted, and tissues were directly incubated with the respective dye, followed by a PBS wash and mounting as described above.

Confocal images were acquired using Zeiss LSM 880 and LSM 980 confocal laser-scanning microscopes. Objective, laser power, and detector gain settings were adjusted according to the experimental requirements and fluorophore intensity. Images were analysed using ImageJ/Fiji (version 2.14.0/1.54p).

### Protein extraction from tissue, Western Blot and Statistics

Early third instar larvae were used to extract total protein by squishing the washed and cleaned larvae with motorised pestle in micro-centrifuge tubes. Standard RIPA lysis buffer (Sigma-Aldrich, R0278) was used after supplementation with additional protease inhibitors (Roche, 04693132001) and 200mM N-ethylmaleimide (NEM; Thermo Fisher Scientific Inc., E3876) (Wang *et al*., 2026). The protein extract was denatured for 10 min at 80°C. Extracted protein was quantified using a standard Bradford Assay using BIO-RAD assay dye reagent (BIO-RAD Laboratories, 5000006) with absorbance being measured at 595 nm on a Genesys^TM^ 10s Vis spectrophotometer.

20ug total protein per loaded well and run in 4-20% BIO-RAD tris-glycine stain free gel (BIO-RAD Laboratories, 4568096) at 150V. Stain free gels were exposed to UV and activated for 45seconds in BIO-RAD ChemiDocMP Gel Imaging System. Western Blot transfer was done using Trans-Blot Turbo Transfer System using Trans-Blot Turbo Transfer packs (BIO-RAD Laboratories, 1704157). Mixed-molecular-weight turbo transfer cycle in 6 minutes. Primary antibody treatment in 5% Tris Buffer Saline Tween-20 (TBST; Scientific Laboratory Supplies Ltd, NAT1246) was done overnight at 4°C and Secondary HRP-conjugated antibody treatment in 5%TBST was done for 2 hours at room temperature, both with gentle agitation. Blots were developed with Amersham™ ECL Select™ (Cytiva, RPN2209) by incubation for 5 minutes at room temperature in dark. Statistical analysis was done using GraphPad prism Version 11.1.0 (218) on data acquired from three independent sets of experiment, n= 3. Early third instar larvae identified from branched but not inverted spiracles, 10 for individual genotype per experiment were pooled into one sample for protein extraction. One-way ANOVA followed by Šídák’s multiple-comparisons test. Significance is indicated as follows: ns (P > 0.05), * (P ≤ 0.05), ** (P ≤ 0.01), *** (P ≤ 0.001).

### Tri-acylgycerol estimation

Triacylglyceride (TAG) levels were calorimetrically from the method previously described by Tennessen et al. 2015 (Lőrincz *et al*., 2017). Third-instar larvae were collected, thoroughly washed and homogenized in 0.05% Tween-20. Samples were then heated at 70°C for 5 minutes to heat inactivate endogenous enzymes. TAG standards and samples were assayed using Triglycerides Liquid reagent (Sentinel Diagnostics/Alpha Laboratories-17624H). Following incubation at 37°C for 10 minutes, absorbance was measured at 546 nm in thermoscientific MULTISKAN skyHigh plate reader. TAG concentrations were calculated from the standard curve and normalized to tissue total protein content measured using Bradford Assay, as described above. Statistical analysis was done using GraphPad prism Version 11.1.0 (218) on data acquired from three independent sets of experiment, n= 3. Early third instar larvae identified from branched but not inverted spiracles, 5 for individual genotype per experiment were pooled into one sample for protein extraction. One-way ANOVA followed by Šídák’s multiple-comparisons test. Significance is indicated as follows: ns (P > 0.05), * (P ≤ 0.05), ** (P ≤ 0.01), *** (P ≤ 0.001).

### Starvation Assay and Drug Treatment

For starvation, early third instar larvae were scooped from rearing tubes, washed thoroughly with water to carefully remove every bit of food. These larvae were incubated on filter paper thoroughly soaked with 20% sucrose (Merck & Co., Inc., 84100) for 4 hours for amino acid starvation and induction of autophagy in fat body cells (Tennessen et. al., 2014).

For drug treatment of live larvae, stock solutions of etoposide (Merck & Co., Inc., 341205) and tunicamycin (Merck & Co., Inc., T7765) were prepared in 100% Dimethyl Sulfoxide (Merck & Co., Inc., 276855) and working concentrations of 100 mM for etoposide and 24 µM for tunicamycin were diluted in an aqueous solution to prepare drug-supplemented fly food. Mid 2^nd^ instar larvae were reared on this drug spiked food for 24 hours. These larvae were hereafter collected from fly food and dissected using the standard protocol as described.

For ex-vivo drug treatment of fat body, larvae were collected and dissected in PBS as per usual. The dissected fat bodies were immediately moved to schneider’s media supplemented with foetus calve serum (10% v/v; Thermo Fisher Scientific Inc.; A5256701) and L-Glutamine-penicillium-streptomycin (1% v/v; Sigma-Aldrich, G6784). This schneider was supplemented with 50 µM for etoposide or 5 µg/ml (6.12 µM) for tunicamycin based on the need of the experiment. The fat bodies were maintained at 25 °C, 70% humidity, 12 h light/12 h dark photoperiod. After the end of treatment period, the fat bodies were washed in PBS, fixed in 4% paraformaldehyde and processed as per usual.

## Results

### Msp300 fragment from Isoform E/G interacts with Atg8a with high confidence

In a previous study conducted in our laboratory (unpublished data), we identified several novel Atg8a-interacting proteins in *Drosophila* through yeast two-hybrid (Y2H) screening. Among the positive interactors, Msp300 fragment (Isoform E and G) emerged as a candidate of particular interest with high confidence in interaction (**Figure. 4**). Msp300 comprises eleven distinct isoforms, which arise from alternative splicing events involving different transcription start site-specifically at exons 1, 3, 18, and 21 as well as various stop and splice sites (Rey *et al*., 2021).

**Figure 1.**
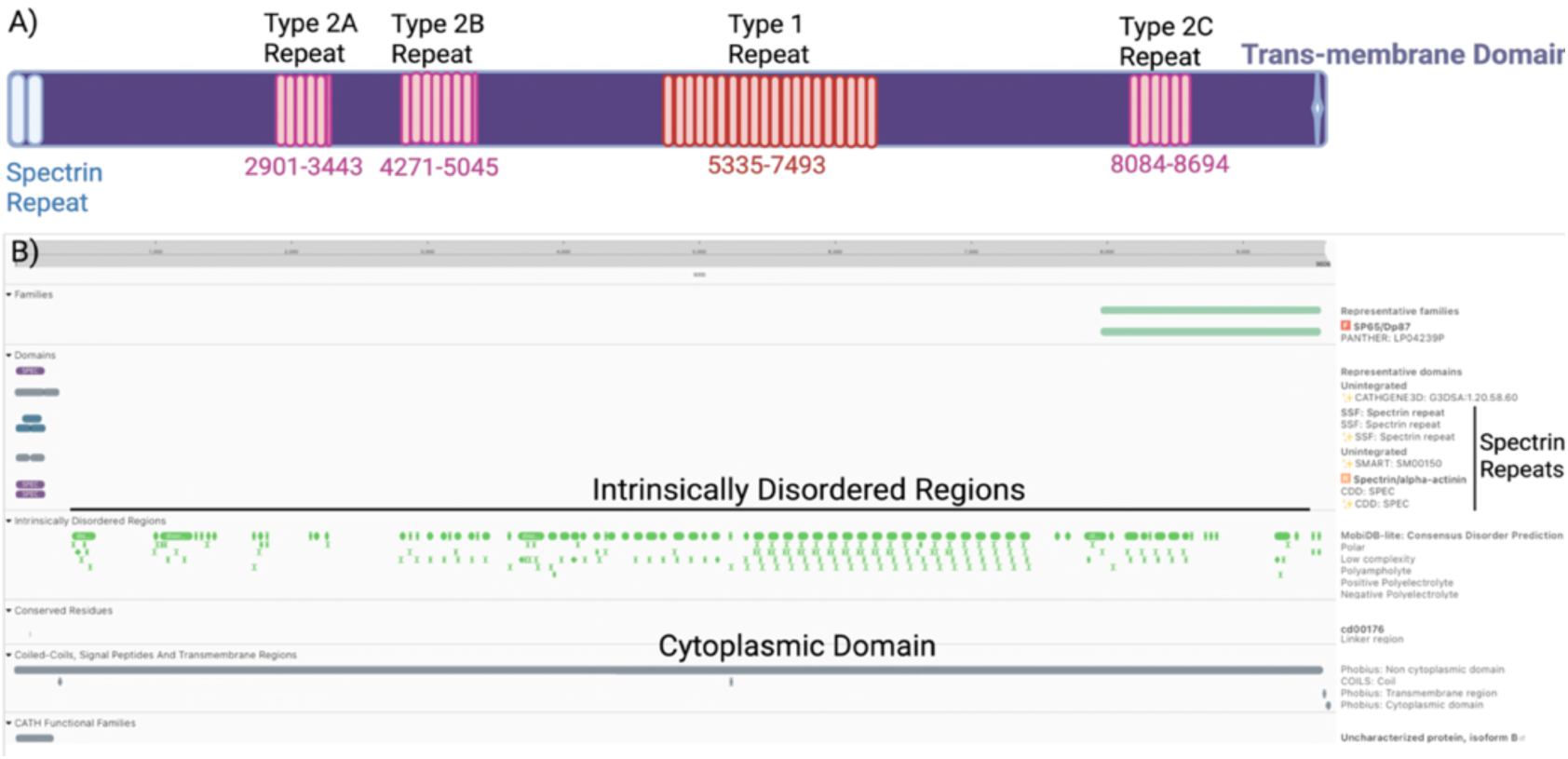
Domain architecture of *Drosophila melanogaster* Msp 300 isoform. **E.** Schematic representation of Msp 300 isoform E. Msp 300 E lacks calponin-homology (CH) and KASH domains and contains two conventional spectrin repeats. The major tandem repeat architecture was determined independently using XSTREAM v1.73, identifying Repeat 2A, Repeat 2B, Type I Repeat and Repeat 2C. Domain annotations were validated using InterProScan, and the C-terminal transmembrane helix was assessed using DeepTMHMM v1.0. Amino-acid coordinates refer to the full-length Msp300-PE sequence. The overall Msp 300 isoform E architecture and repeat nomenclature are consistent with the published characterization Msp 300 isoform by Morton *et al*. (2025). Created in https://BioRender.com

**Figure 2.**
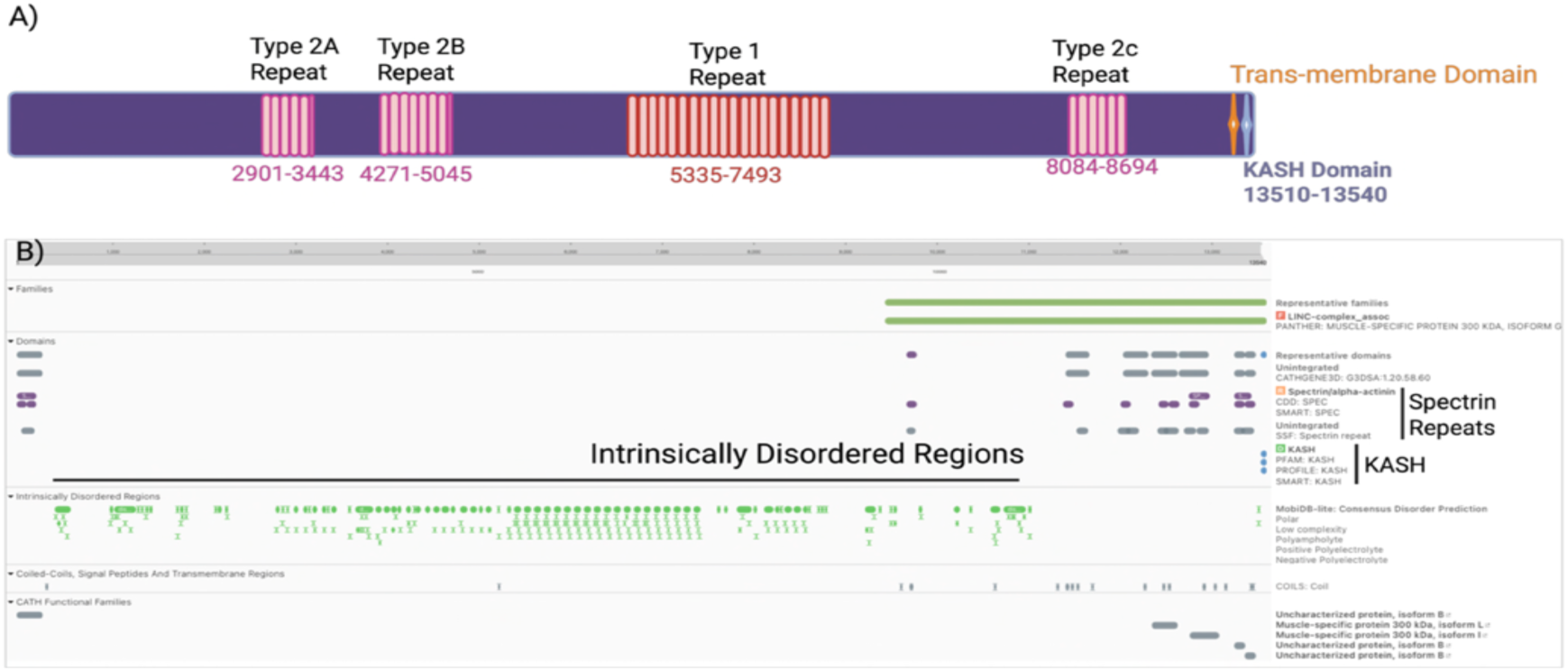
Sequence based domain architecture of *Drosophila melanogaster* Msp 300 isoform. **G.** Msp300 isoform G amino acid sequence was analyzed using XSTREAM v1.73, InterProScan, and DeepTMHMM. XSTREAM identified four major tandem-repeat blocks: Repeat 2A, Repeat 2B, Repeat 1, and Repeat 2C. InterProScan identified spectrin-repeat and KASH annotations, while DeepTMHMM predicted an additional C-terminal transmembrane helix at aa 13488–13509. Msp 300 G has been reported to contain 21 spectrin repeats and no CH domains. Amino-acid coordinates refer to the full-length Msp300-PG sequence. Created in https://BioRender.com

**Figure 3.**
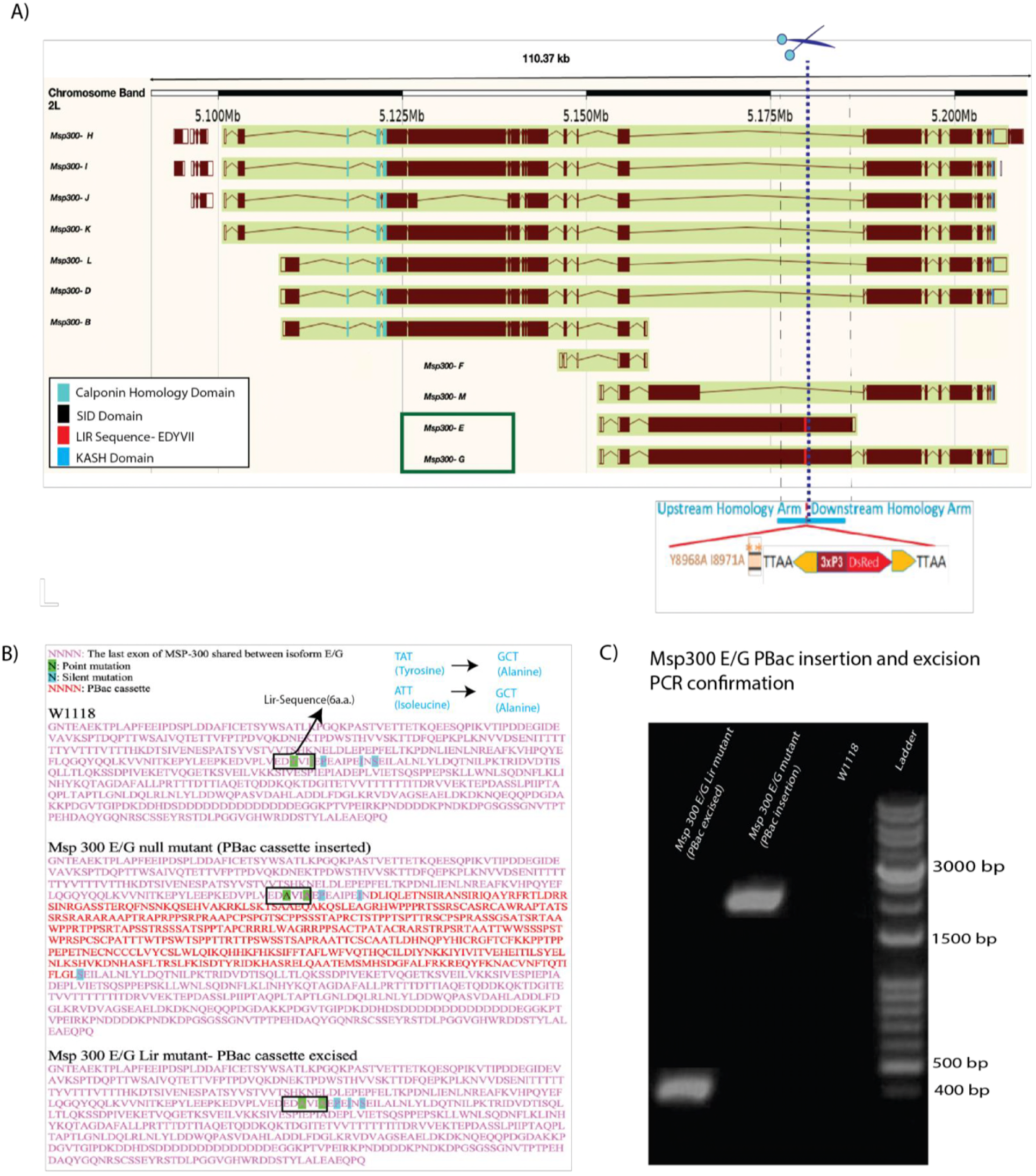
Generation of Msp300-E and Msp300-G mutant flies by PBac-mediated gene disruption and LIR-sequence point mutation. **(A)** Schematic of CRISPR/Cas9-mediated genome editing by homology-directed repair using the Pbac-DsRed knock-in cassette (WellGenetics Inc., Taipei, Taiwan). **(B)** Genetic manipulation of the Y8968A and I8971A residues in Msp300-E downstream of the 6-aa LIR sequence. Insertion of the PBac cassette in Msp300-E mutants introduces point mutations in the LIR sequence and results in loss of both Msp300-E and Msp300-G isoforms due to their shared exon. In the Msp300-E/G LIR mutant, the PBac cassette is excised, leaving only the two-amino-acid point mutation in the LIR sequence. **(C)** PCR confirmation of PBac cassette insertion and excision in Msp300-E/G mutants.

**Figure 4.**
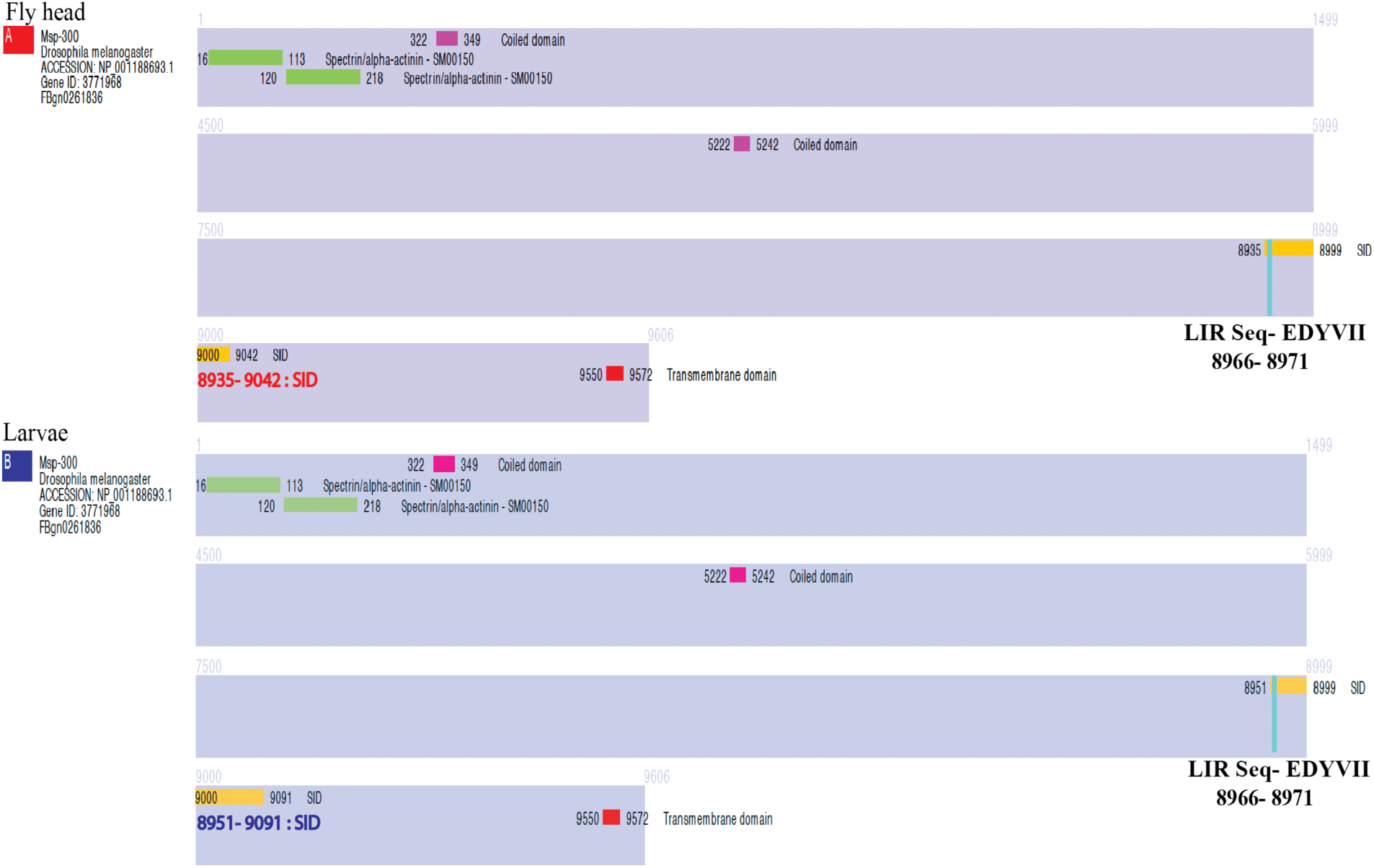
High-throughput yeast two-hybrid screening in Drosophila identifies Msp300 as a novel putative Atg8a interactor. High-throughput yeast two-hybrid screening (Hybrigenics Services, Paris, France) identified Drosophila Msp300-E as an Atg8a-interacting protein, with a Predicted Biological Score (PBS) of A (very high confidence) in fly heads and B (high confidence) in larvae. An iLIR software-predicted LIR interaction motif (blue box) was identified within the selected interaction domain (yellow box) present in the Msp300 fragments (purple box) that interacted with Atg8a.

Notably, Isoforms E and G share a portion of exon 18, a feature absent in all other isoforms. In this Y2H analysis we specifically observed the interaction between Msp300 Isoform E and G with Atg8a and established that this interaction was mediated by a LIR motif (6aa-EDVYII) which was found to be present in cytoplasmic region of Msp300 isoform (E/G) and is important feature of autophagy receptors (**Figure. 5**).

**Figure 5.**
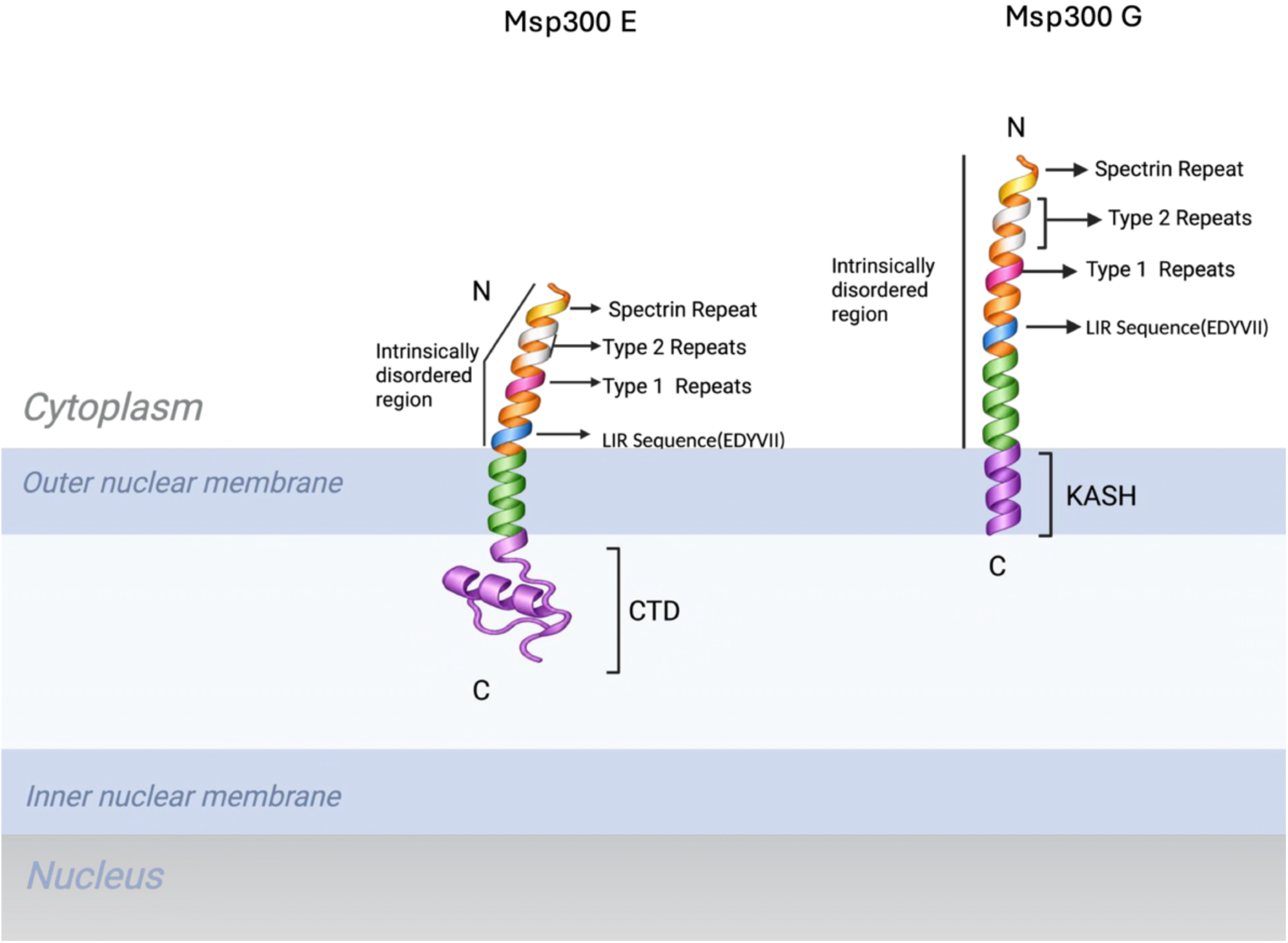
Illustration of the topology of *Drosophila melanogaster* Msp300 isoform E and G at the outer nuclear membrane. Schematic model of Msp300 E and G at the outer nuclear membrane, based on the domain organisation and structural predictions reported by Morton et al. (2025). Msp300 isoforms E and G are depicted as having a large N-terminal cytoplasmic region and a C-terminal transmembrane domain anchoring the proteins in the outer nuclear membrane. Msp300 has a KASH domain that attributes ONM localisation, while Msp300 has a predicted transmembrane domain and a 35-aa C-terminal domain (CTD), which is predicted to reside within the perinuclear space. The previously described spectrin repeat and the Type 1 and Type 2 tandem-repeat regions identified using XSTREAM v1.73 are indicated, together with the six-aa EDYVII LIR sequence predicted using iLIR. Created in BioRender.com.

We performed targeted Y2H assays using a Selected Interaction Domain (SID) (8909-9217) fragment of both Msp300 isoform E as well as G, identified from previous untargeted Y2H. In this study, we observed a robust interaction between Atg8a and the Msp300 E SID (8909-9217) fragment. For the Msp300 E interaction experiment, SMAD and SMURF protein interaction was used as a positive control. For the negative control, Atg8a, Msp300 E SID, Msp300 E SID deleted, and Msp300 E LIR deleted fragments were tested against empty vector to eliminate possibility of autoactivation. No colonies were observed in this case on DO3 media lacking histidine. However, in case of interaction between Atg8a and Msp300 E SID fragment colonies were observed in DO3 media lacking histidine. The interaction between prey and bait fragment drives production of histidine which helps in survival on histidine deficient media. Notably, deletion of the SID fragment abolished this interaction, confirming its necessity for binding.

Most interestingly, upon mutation in 2aa within the 6aa from LIR domain (EDYVII → EDAVIA) in SID fragment this interaction is lost. This strongly suggested that the interaction between Msp300 E and Atg8a is LIR domain mediated (**Figure. 6**).

**Figure 6.**
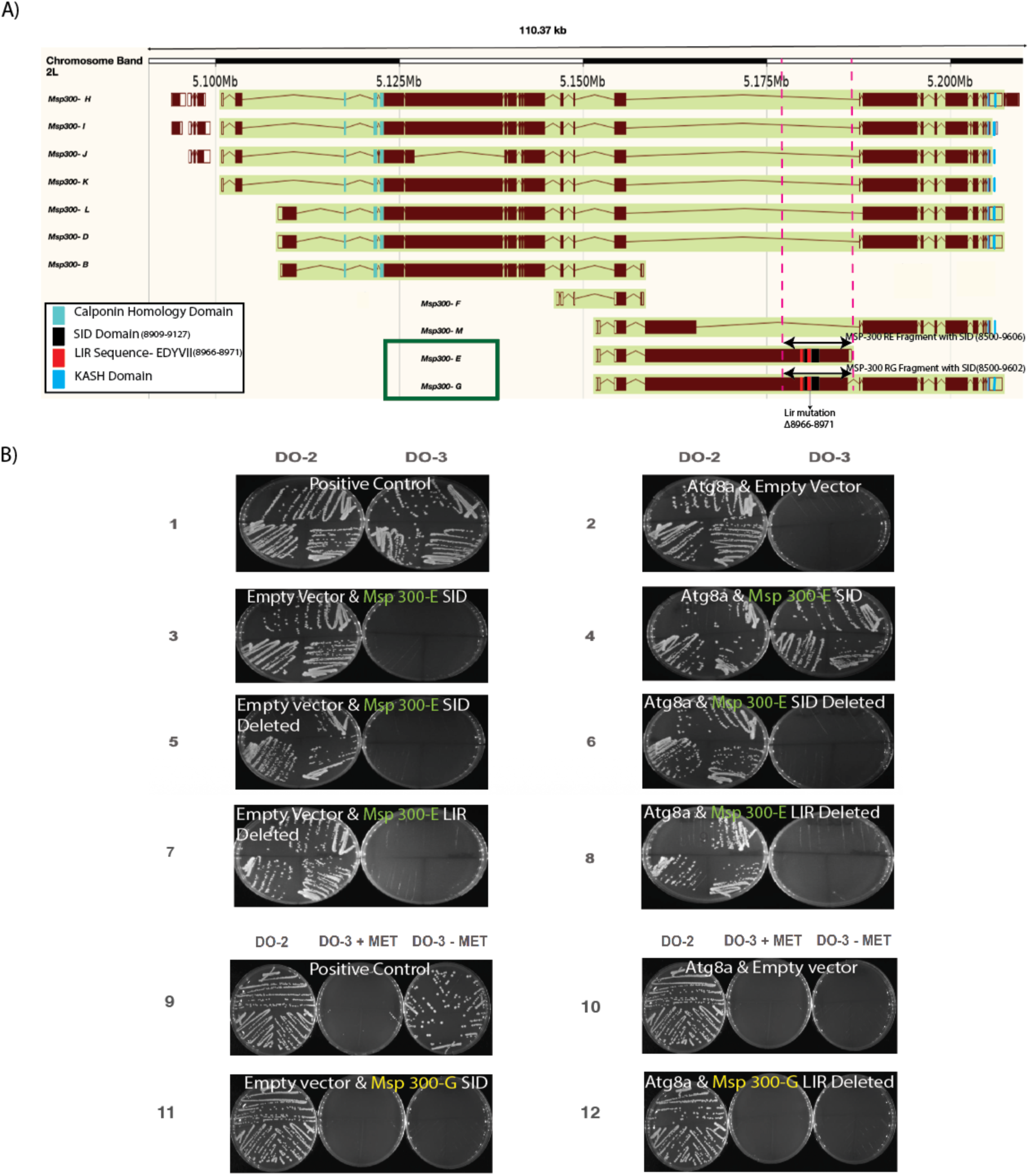
Yeast two-hybrid (Y2H) mutation analysis confirms Msp300 and Atg8a interaction is LIR mediated. **(A)** Schematic representation of Msp300 isoforms highlighting key domains and fragments used in Y2H assays. Isoform E contains a Selected Interaction Domain (SID; residues 8909–9127) that initially demonstrated interaction with Atg8a. Test fragments encompassing residues 8500– 9606 (isoform E) and 8500–9602 (isoform G) were subsequently analyzed for Atg8a binding. **(B)** Results from yeast two-hybrid (Y2H) interaction studies (Hybrigenics Services, Paris, France) confirmed that Atg8a interacts with MSP300 test fragments. Deletion of the selected interaction domain (SID) abolished this interaction, demonstrating that the SID is required for Atg8a binding. Further mutational analysis showed that deletion of the 6-amino-acid LC3-interacting region (LIR) motif within the SID also abolished the interaction. These results indicate that the LIR motif within the SID is essential for Atg8a-MSP300 binding.

For Msp300 G, the first exon used for alternative splicing is same as Msp300 E and the protein sequence of SID fragment tested for Msp300 E is the same as Msp300 G, therefore we only tested interaction between Msp300 G SID fragment-LIR deleted with Atg8a. In this case, internal inducible control bait and prey fragment interaction was tested in DO2, DO3 + Met and DO3 - Met media. Colonies were observed in all the three systems confirming the reporter activation, selection conditions and detection of true interaction. For the negative controls, Atg8a and Msp300 G SID fragment LIR deleted were individually checked for interaction with empty vector and no growth was observed in DO3 + Met and DO3 - Met media but growth of colonies was observed in DO2 media indicating yeast is viable and has relevant plasmids and even so there is no interaction between the bait and the pray fragment. In case of the interaction between Msp300 G SID fragment LIR deleted and Atg8a fragment, similar pattern was observed with no growth in DO3 + Met and DO3 - Met media but growth of colonies in DO2 media (**Figure. 6**). This Y2H mutational analysis study confirmed Msp300 E/G is putatively a strong interactor of Atg8a and, this Msp300 E/G and Ag8a interaction is LIR domain mediated.

We also performed AlphaFold analysis to test in-silico interaction between Msp300 isoform E and G. For Msp300 isoform E with 9606aa, aa sequence was broken into 1500aa long overlapping 7 fragments. These fragments were then subsequently individually tested for interaction with Atg8a. The 6 fragments gave ipTM values ranging from 0.19-0.85 and no protein-protein interaction was observed within these. However, in the last fragment (residues 8107-9606aa) gave an ipTM of 0.95; piCS 0.86; AlphaBridge 0.98 and within this fragment we identified a single interface in which a 17-residue stretch (residues 8967-8983aa) docked onto Atg8a, 44-55aa and 59-63aa (**Figure. 7**). Similarly, in Msp300 isoform G, 9 fragments gave ipTM values ranging from 0.18-0.79 no protein-protein interaction was observed within these. But as expected due to exon sharing, one fragment (residues 8035-9523aa) gave an ipTM of 0.95(Scores: ipTM 0.95; piCS 0.86; AlphaBridge 0.98 and single interface that we found in Msp300 isoform E, 17 residue stretch (residues 8967-8983aa) showed interaction with Atg8a (**Figure. 8**). Therefore, for control experiments only Msp300 isoform E was further tested.

**Figure 7.**
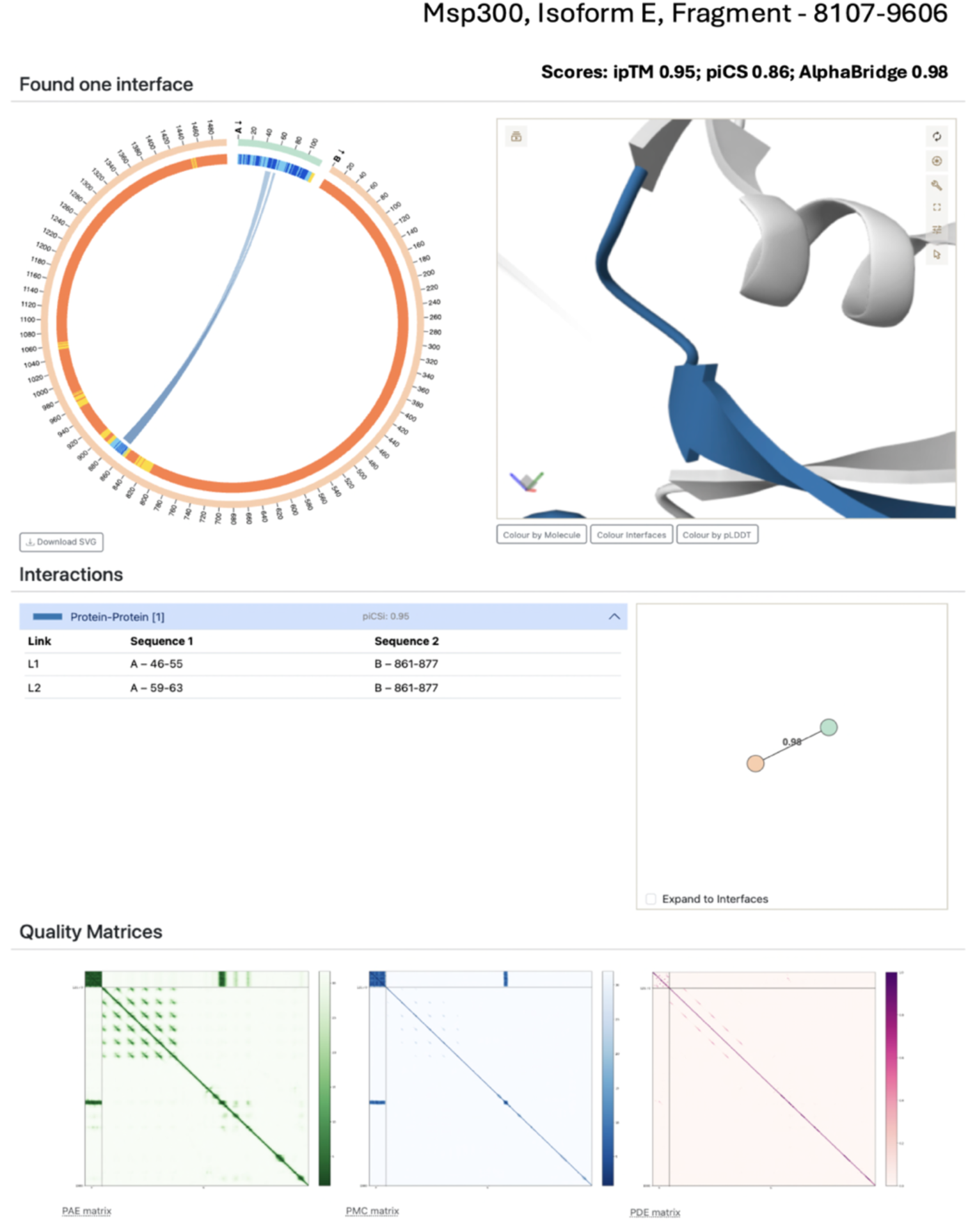
AlphaFold 3 prediction of the interaction between the 7^th^ fragment (8107-9606 aa) of Msp300 isoform E and Atg8a. The predicted complex identified a candidate interaction interface within the 46-55aa in Atg8a and 8967-8983aa in Msp300 isoform E, with high ipTM of 0.95 and AlphaBridge score of 0.98.

**Figure 8.**
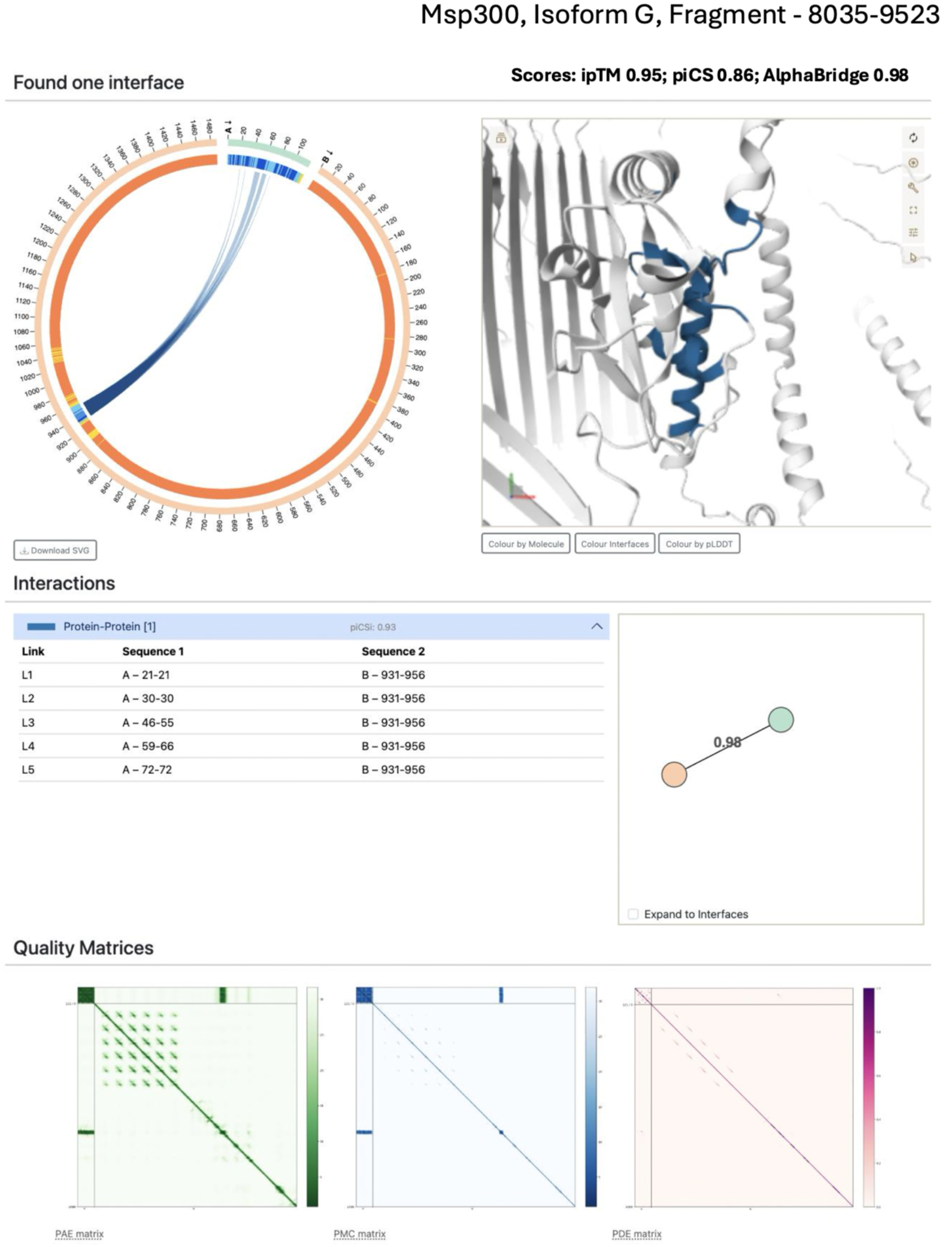
AlphaFold 3 prediction of the interaction between the 7^th^ fragment (8035-9523 aa) of Msp300 isoform G and Atg8a. The predicted complex identified a candidate interaction interface within the 46-55aa in Atg8a and 8967-8983aa in Msp300 isoform E, with high ipTM of 0.95 and AlphaBridge score of 0.98.

Because AlphaFold can return confident predictions that are not true interactions, specially in disordered regions, we additionally tested with different fragment lengths, mutated interaction motif and tested unrelated protein with similar disordered region. The fragment that showed interaction was trimmed to 140 residues around the site of interaction to eliminate effect of disordered region; the interaction persisted (ipTM 0.76; piCS 0.39; AlphaBridge 0.90) (**Figure. 9**). A scrambled fragment with only region of interaction shuffled while preserving its composition and flanking sequence, the ipTM fell to 0.28 (Scores: ipTM 0.28; piCS 0.00; AlphaBridge 0) (**Figure. 10**). We also tested Atg8a with LIR mutated Msp300 E fragment (EDYVII to EDAVIA) and interestingly found that interaction was still there (Scores: ipTM 0.83; piCS 0.56; AlphaBridge 0.79) and localised close to same residue 46^th^ and 50^th^ aa interacted individually with 8960-68962aa (**Figure. 11**). Pairing Atg8a with full length unrelated protein with high intrinsically disordered region-no interaction was predicted (ipTM 0.45; piCS 0.21; AlphaBridge 0.44) (**Figure. 12**). Together these results are consistent with a sequence-specific, motif-mediated interaction between Msp300 and Atg8a.

**Figure 9.**
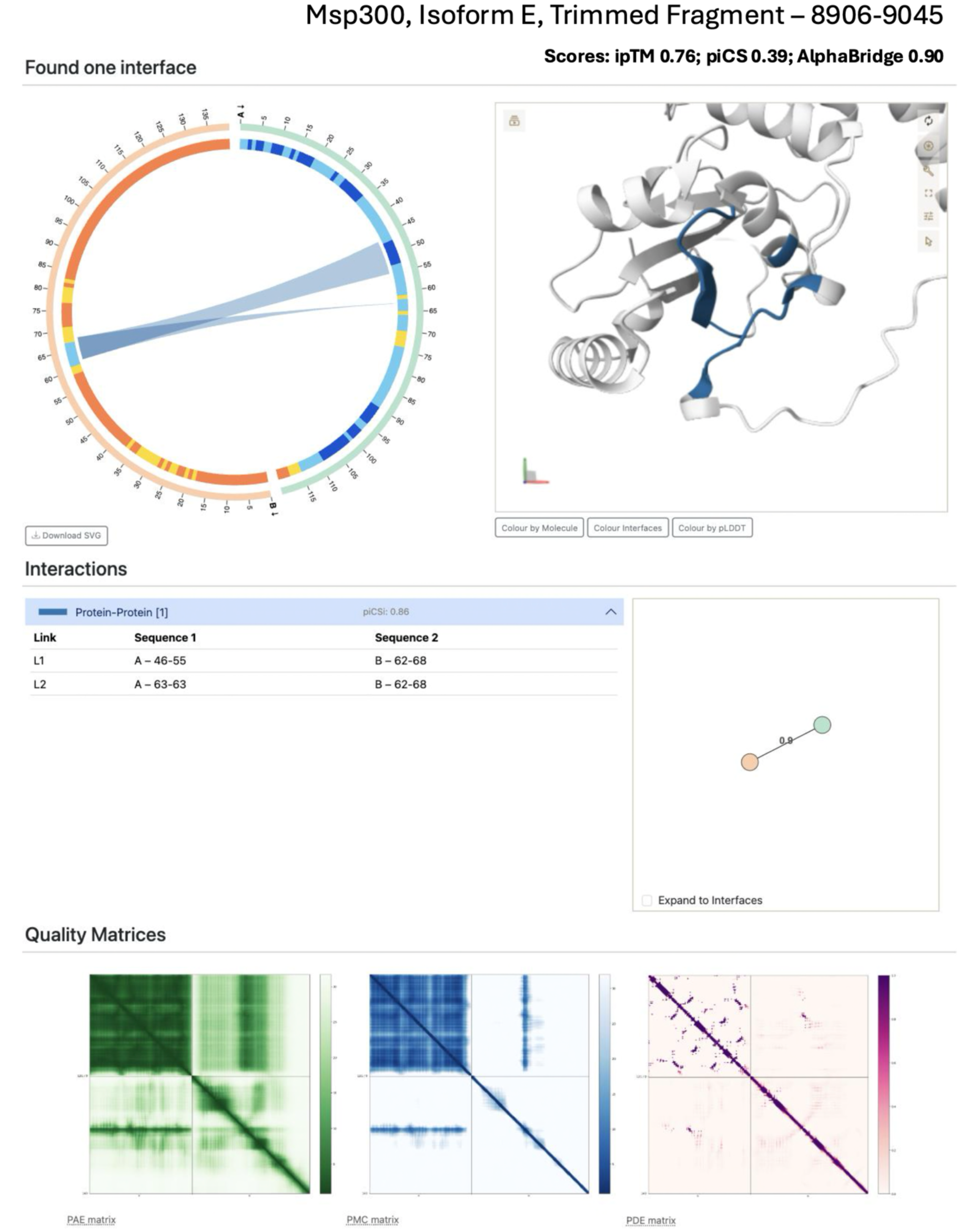
AlphaBridge analysis of the trimmed 140-amino-acid target fragment with Atg8a. The predicted interface was retained in the trimmed construct, further supporting the idea that interaction is from sequence of interest and not flaking sequence of the original fragment.

**Figure 10.**
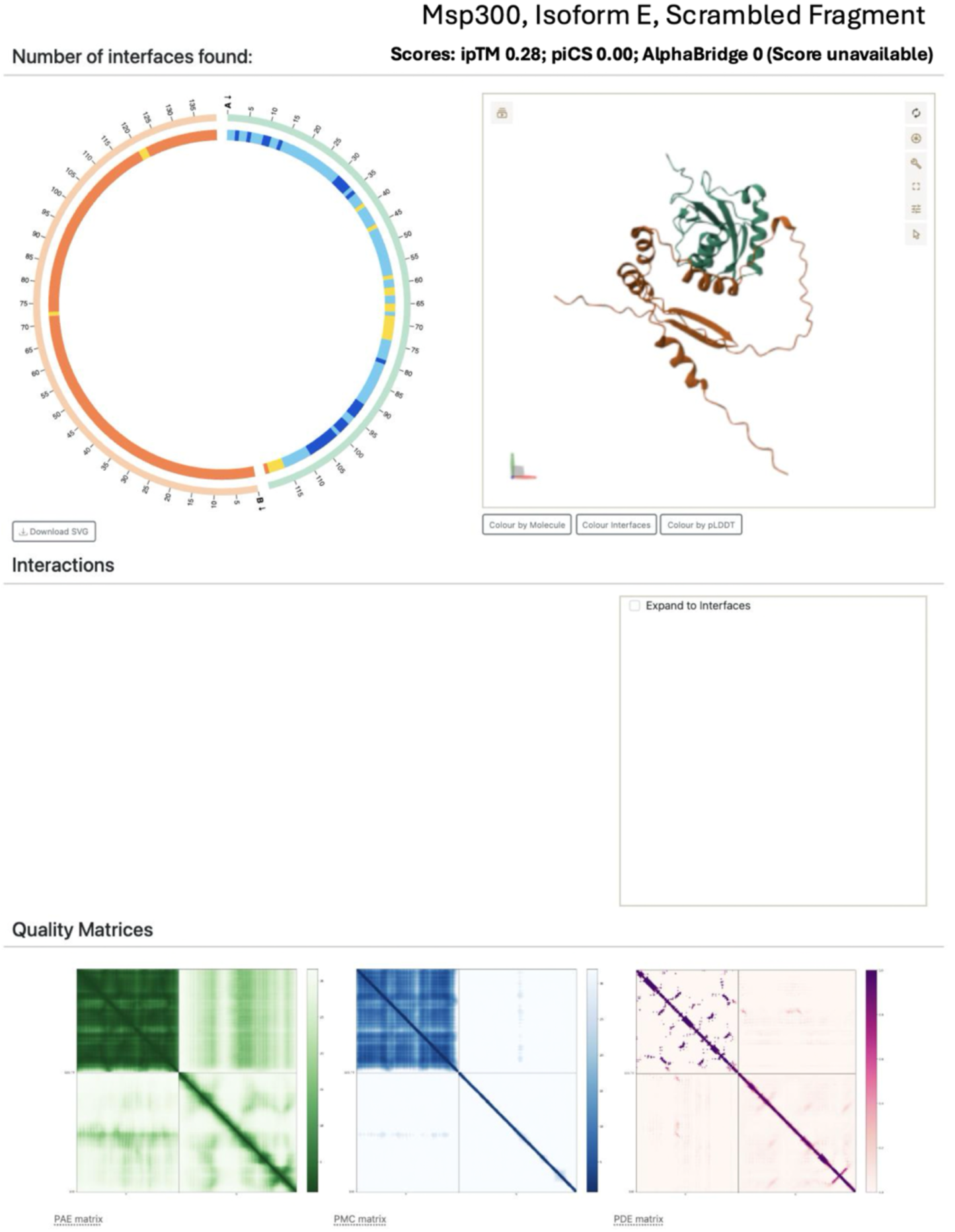
AlphaBridge analysis of the scrambled-fragment control with Atg8a. The model showed substantially lower interface scores and no detected interface compared with the native fragment confirming the necessity of sequence for the interaction.

**Figure 11.**
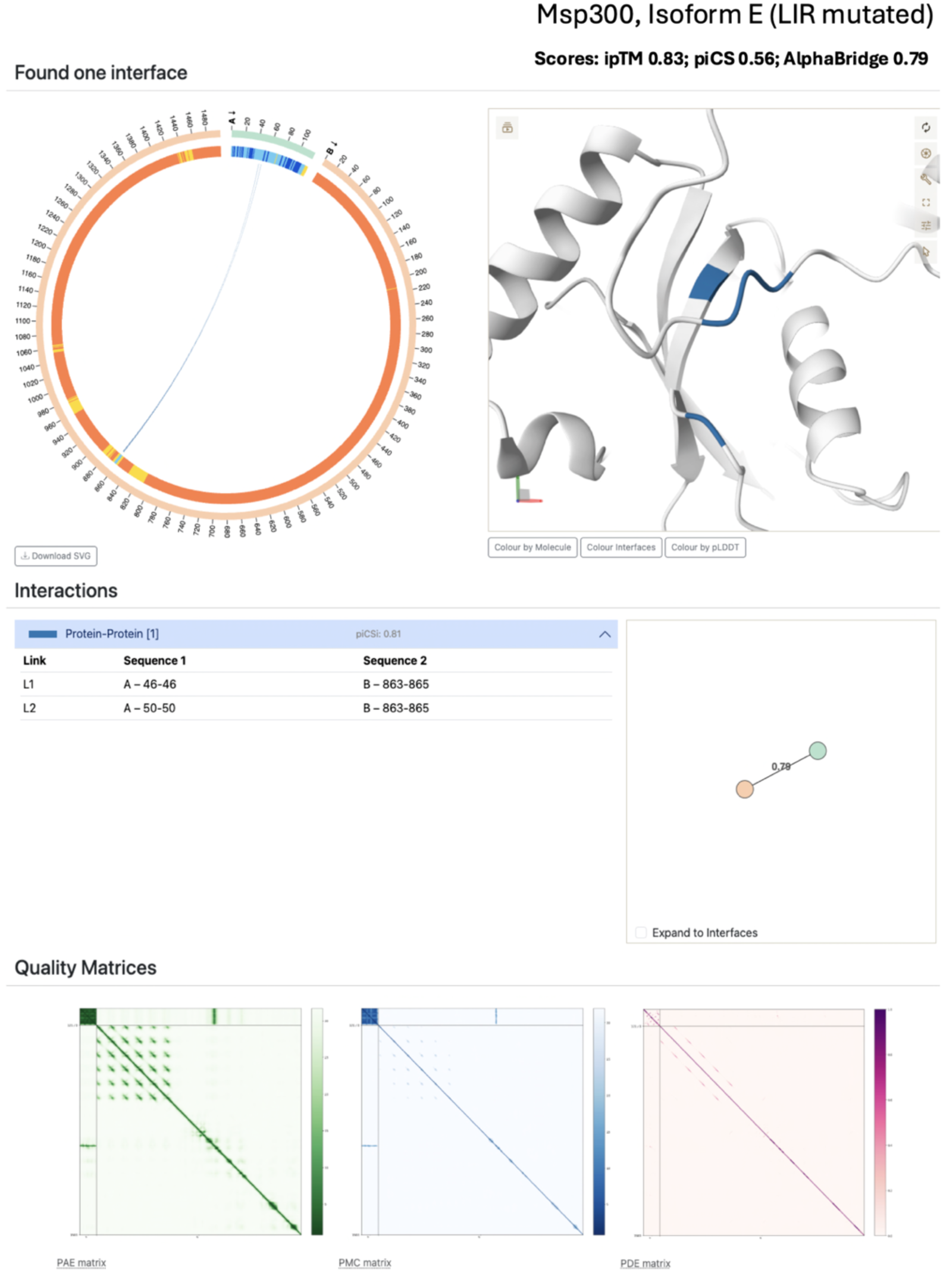
AlphaBridge analysis of the mutated target fragment with Atg8a. The model retained a predicted interface, although the interface scores were reduced relative to the original construct. Potentially defining the necessity of flanking 2 amino acid to form a stable interaction between the two fragments.

**Figure 12.**
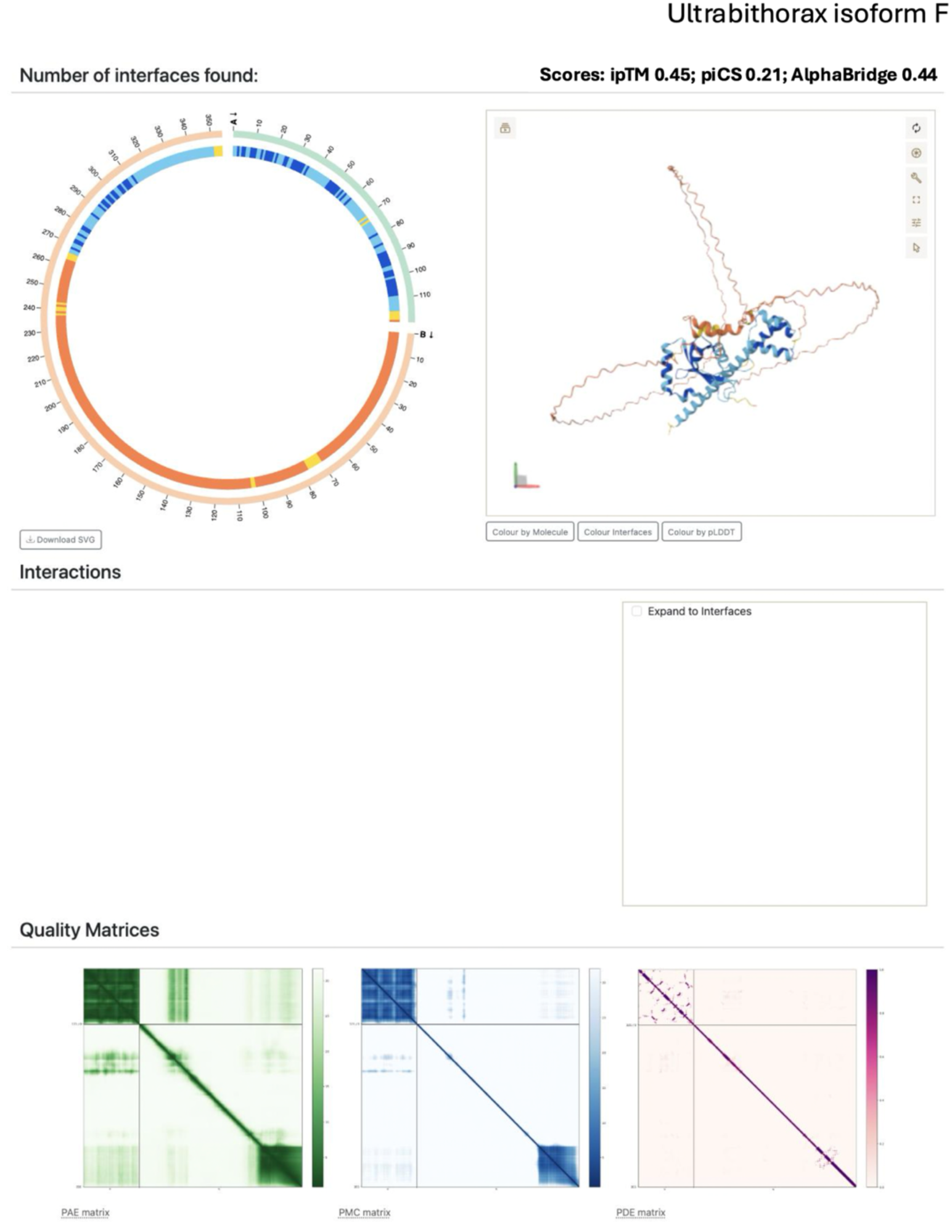
Structural visualisation of the predicted protein-protein interface. The model highlights the relative arrangement of Atg8a and the target fragment, with candidate inter-residue contacts indicated in the structure.

### Impacted nuclear positioning in Msp300 and Atg8a mutant

Given the observed interaction between Atg8a and Msp300, and the role of Msp300 in organising the cytoskeleton and nucleoskeleton, as well as in nuclear positioning, we examined whether Atg8a interacts with Msp300 to facilitate nuclear circularity and centrality. In the fat body cells, Msp300 and Atg8a mutants were checked for their requirement to position the nucleus at the centroid relative to F-actin which delineated the cell boundary. In body wall muscles of 3^rd^ instar larvae of W118 isogenic control lines, nucleus is evenly spaced out and in fat body it is distinctly circular and located at the centre of the cell (**Figure. 13 A-A’**). In Msp300 complete clonal mutant (RFP -ve cells deficient in all Msp300 isoforms) in the body wall muscle we saw nucleus clustering phenotype and in RFP -ve fat body cell’s nucleus was pushed to the corner and appeared collapsed and non-circular **(Figure. 13 B-B’).** In the body walls muscle of Msp300 E/G null mutant similar nuclear clustering was observed and in fat body cells all the nuclear appeared displaced from the centre and did not look circular **(Figure. 13 C-C’).**

**Figure 13.**
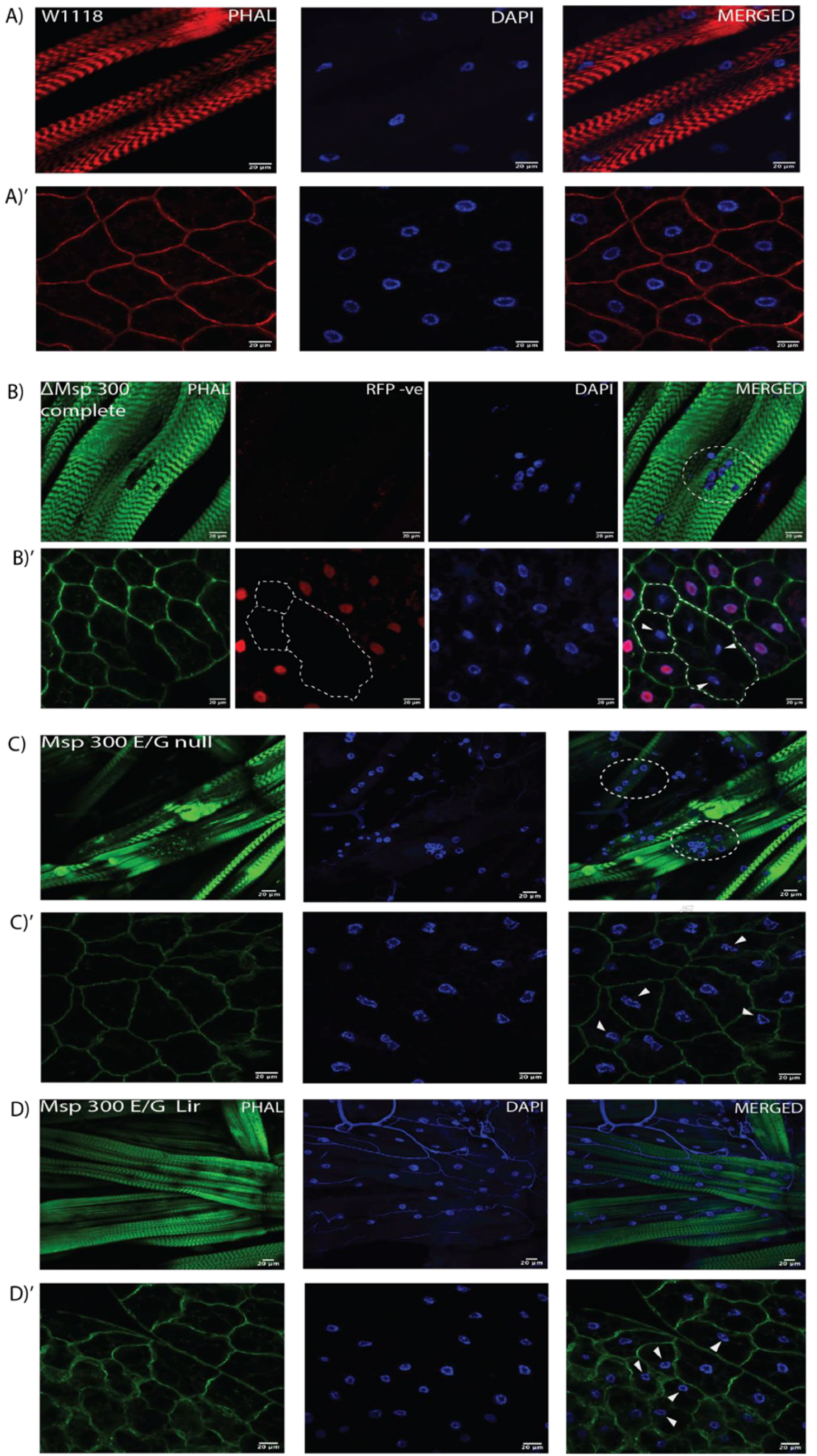
Nuclear organization in larval body wall muscle and fat body cells of W118 and Msp300 mutants. (A′–D′) Representative confocal images of larval body wall muscle from W1118, Msp300 complete clonal, Msp300 E/G null, Msp300 E/G LIR mutants. **(A–D)** Representative confocal images of larval fat body cells from W1118, Msp300 complete clon Msp300 E/G null, and Msp300 E/G LIR mutants. **(A, A′)** In W1118, nuclei in body wall muscle are evenly distributed, wherea nuclei in fat body cells are circular and centrally positioned. **(B, B′)** In Msp300 complete clonal body wall muscle and fat body cells, RFP-negative cells show altered nuclear organization. In body wall muscle, nuclei lose their even distribution and cluste together (white circle). In fat body cells, nuclei appear compressed and displaced towards one corner of the cell. **(C, D)** Body muscle from Msp300 E/G null and Msp300 E/G LIR mutants, respectively. Msp300 E/G null mutants show displaced (non-central) and irregularly shaped nuclei (white circle), whereas Msp300 E/G LIR mutants show nuclear arrangement and morphology resembling those of W1118. **(C′, D′)** Fat body cells from Msp300 E/G null and Msp300 E/G LIR mutants, respectively. In both mutants, nuclei are non-centrally positioned (white arrows), with an apparent alteration in nuclear circula

Interestingly in Msp300 E/G LIR mutant however, in the body wall muscles the nucleus was evenly spaced out like the control lines while in the fat body like Msp300 E/G null mutant nucleus was pushed to the corner and did not appear perfectly circular **(Figure. 14 A-B).**

**Figure 14.**
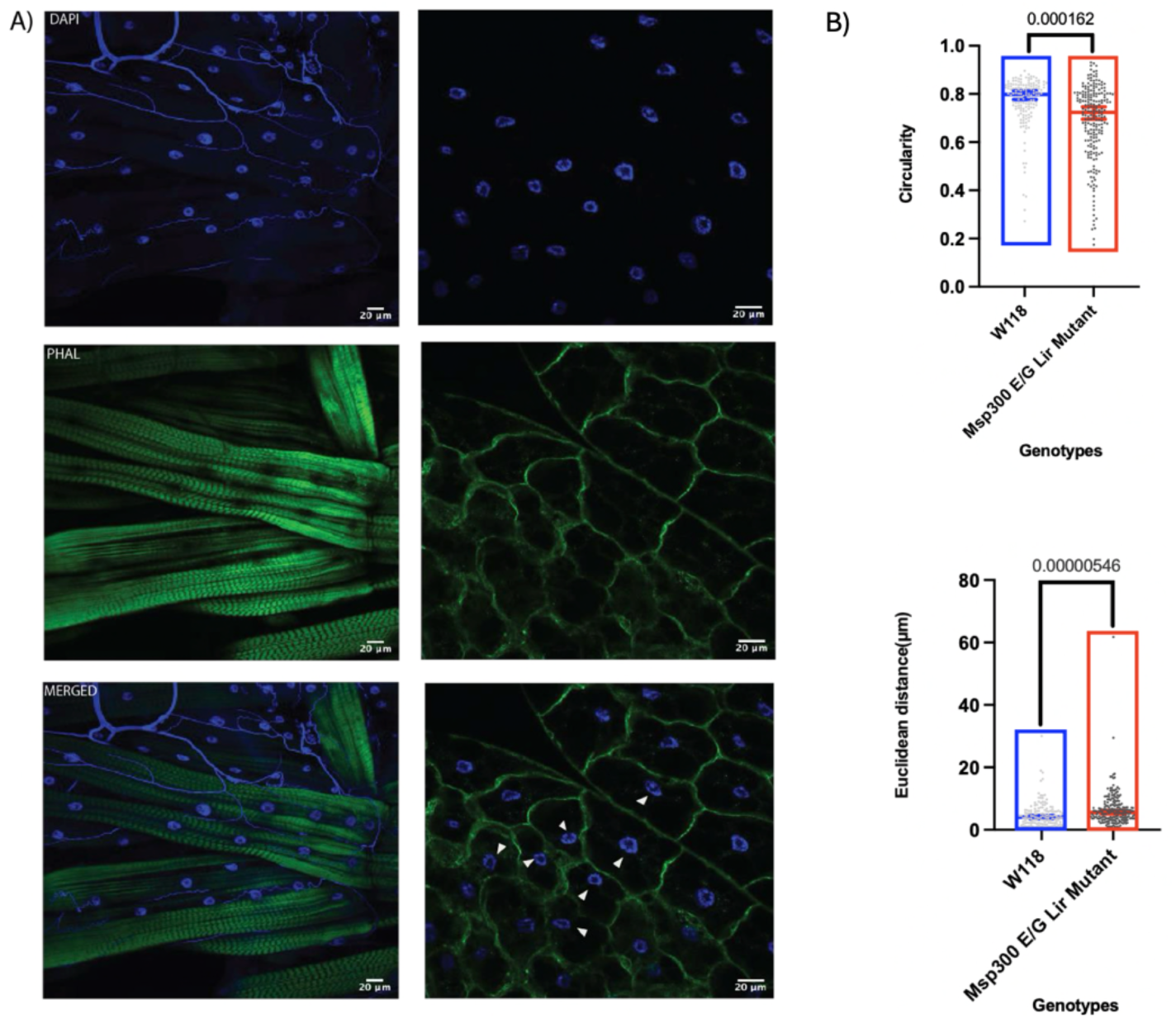
Role of the LIR motif in Msp300 isoforms E and G for maintaining nuclear shape and positioning in the fat body of second-instar larvae. **(A)** Representative confocal images showing nuclei in fat body cells of Msp300 E/G LIR mutants compared to wild-type (WT) controls. Mutant cells exhibit displaced (non-centric) and irregularly shaped (non-circular) nuclei, unlike the centrally positioned and circular nuclei observed in WT cells. F-actin is visualized in green (phalloidin staining), and nuclei are labeled in blue (DAPI). White arrows point to displaced nuclei and non-circular nuclei. **(B)** Quantification of nuclear circularity and centrality. Data are presented as mean ± SD. Statistical analysis was performed using a non-parametric Mann–Whitney test on Euclidean distance (for centrality) and circularity measurements (120 ≤ n ≤ 214). Significance is indicated as follows: ns (P > 0.05), * (P ≤ 0.05), ** (P ≤ 0.01), *** (P ≤ 0.001).

In Atg8a RNAi clonal mutants (GFP +ve cells representing Atg8a mutant cells) we checked for nucleus circularity and centrality (**Figure. 15A**). The RNAi knockdown was confirmed by checking accumulation of Ref (2)P, a substrate of autophagy which is cleared timely if autophagy is functional. Atg8a RNAi clonal GFP +ve cells accumulated significantly more Ref (2)P than wild type GFP -ve cells (**Figure. 15 B**). Immunofluorescence analysis revealed that in Atg8a clonal RNAi mutants, the centrality of nucleus was disturbed, and the shape of nuclei was also not quite circular which suggested a supporting role of Atg8a in Msp300 mediated positioning of the nuclei.

**Figure 15.**
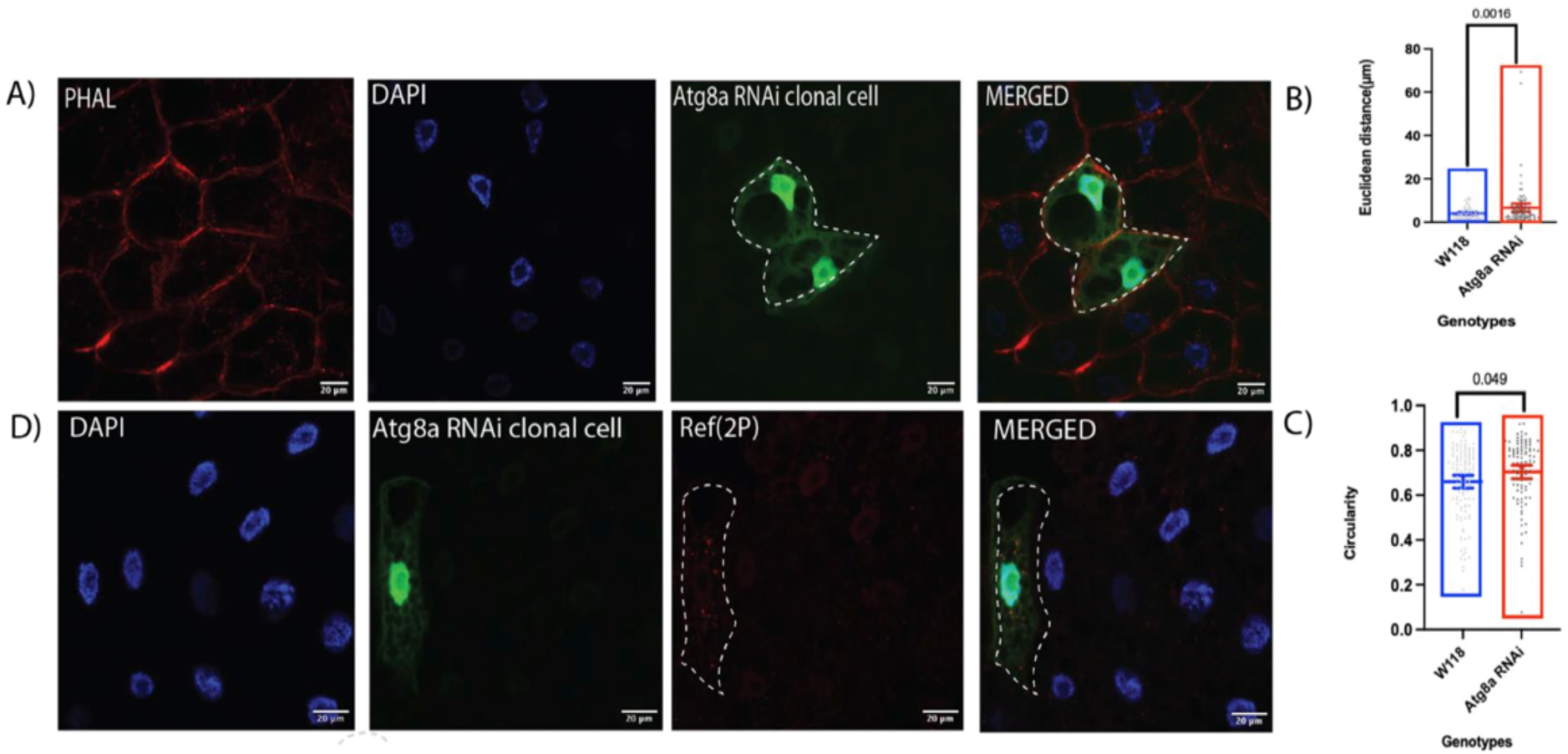
Clonal analysis of Atg8a RNAi mutants for nuclear shape and positioning in the fat body cells of second-instar larvae. **(A**) Representative confocal images of fat body cells showing clonal Atg8a RNAi mutant cells (outlined in white) surrounded by wild-type (WT) cells. F-actin is visualized in red (phalloidin staining), nuclei are labeled in blue (DAPI), and GFP marks Atg8a RNAi mutant clones (green). Mutant cells exhibit displaced (non-centric) and irregularly shaped (non-circular) nuclei compared to the centrally positioned and circular nuclei in WT cells. **(B-C)** Quantification of nuclear circularity and centrality in WT and RNAi cells. Each point represents an individual cell from fat bodies collected from 10 flies per experiment (four independent experiments); bars indicate mean ± 95% CI. Circularity was analyzed using a linear mixed-effects model, and Euclidean distance using linear regression on log-transformed values. n = 221 cells. **(D)** Confirmation of Atg8a RNAi knockdown by assessing Ref (2)P protein accumulation.

### Lamin localisation defects in the Msp300 and Atg8a mutants

In *Drosophila*, there are two lamin genes-the B-type (Lamin Dm0) and the A-type (Lamin C) of which A-type Lamins are rigid and responsible for mechanostress absorption and structural reinforcement by attributing nuclear stiffens while the B-type Lamins are more flexible and support nuclear envelope integrity and restore nuclear shape after mechanostress (Evangelisti *et al*., 2022). Since we observed nuclear displacement and shape defects, we wanted to check integrity of nucleoskeleton in Msp300 and Atg8a mutants. In control line cells, there was a neat ring of both Lamin B and Lamin C outside the nucleus (**Figure. 16 B and 17 A**). In Msp300 complete clonal mutant (RFP−ve cells), Lamin C as well as Lamin B were disintegrated, and cytoplasmic blobs of Lamin B were observed (**Figure. 16 A and 17 B**). In Msp300 E/G null mutant similarly, Lamin C and Lamin B were perforated and disintegrated. The nucleic acid protruded out of the Lamin B ring, indicating a deformed nucleus (**Figure. 16 C and 17 C**). In Msp300 E/G LIR mutant, Lamin C and Lamin B were both perforated and disintegrated, with nucleic acid blebbing out of the Lamin B and Lamin C rings, indicating an impacted nucleus (**Figure. 16 D and 17 D**). In Atg8a RNAi clone, Lamin C was perforated and patchy, while Lamin B although intact, showed nucleic acid spillage outside the Lamin ring (**Figure. 16 B and 17 E**).

**Figure 16.**
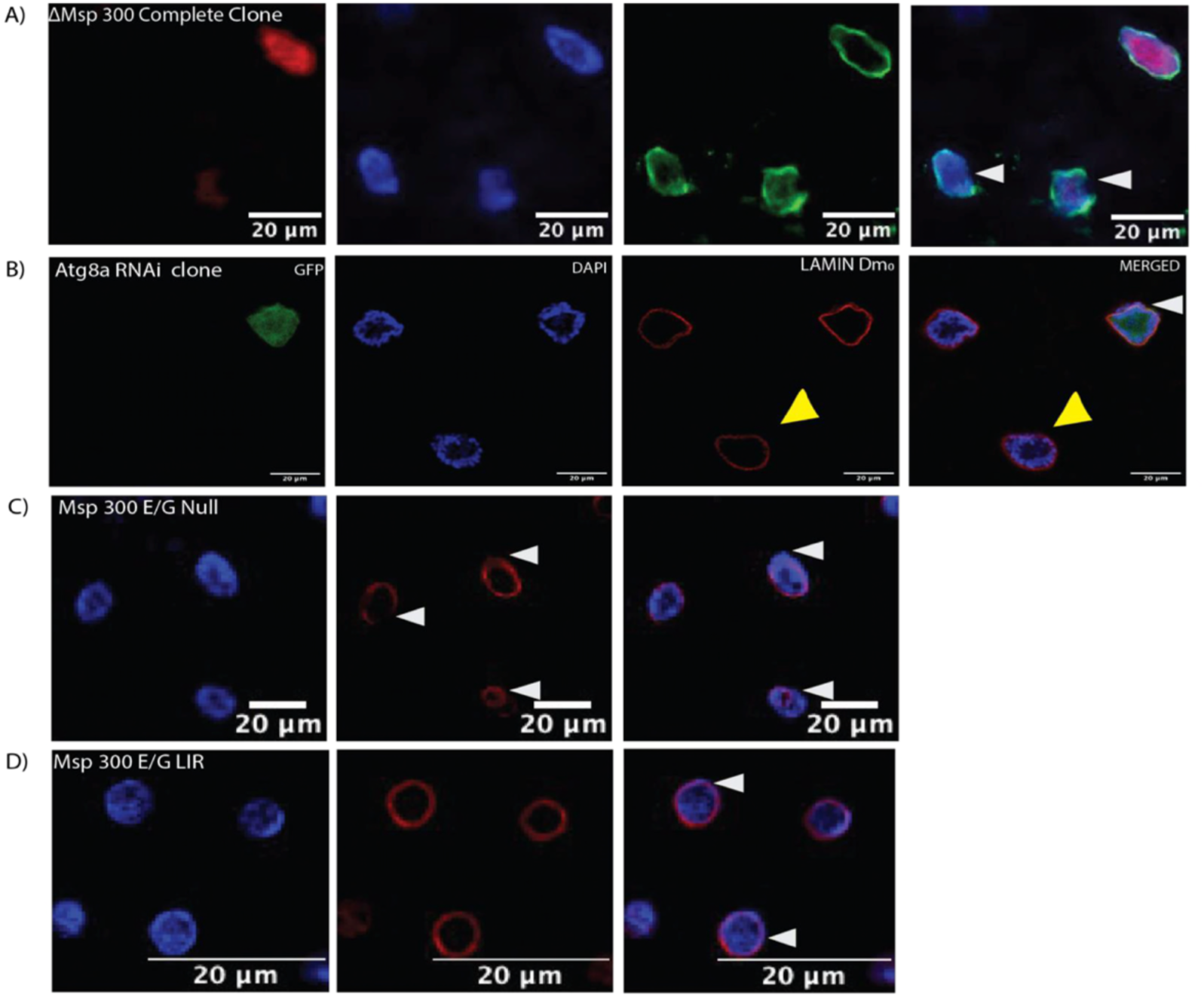
Disrupted Lamin Dm0 organization in Atg8a and Msp300 mutants. Representative images of Lamin Dm0 organization in fat body cells from late second-instar larvae. In wild-type cells, Lamin Dm0 surrounds the nucleus, with the nucleus enclosed within the Lamin Dm0 network (yellow arrow). Msp300-null clonal cells, identified as RFP-negative cells using the FLP-out system, show irregular and disrupted Lamin Dm0 organization. In Atg8a RNAi GFP-positive clonal cells and Msp300 E/G LIR mutants, the nucleus extends beyond the Lamin Dm0 boundary. In Msp300 E/G null mutant, Lamin Dm0 appears deformed with areas of accumulation.

**Figure 17.**
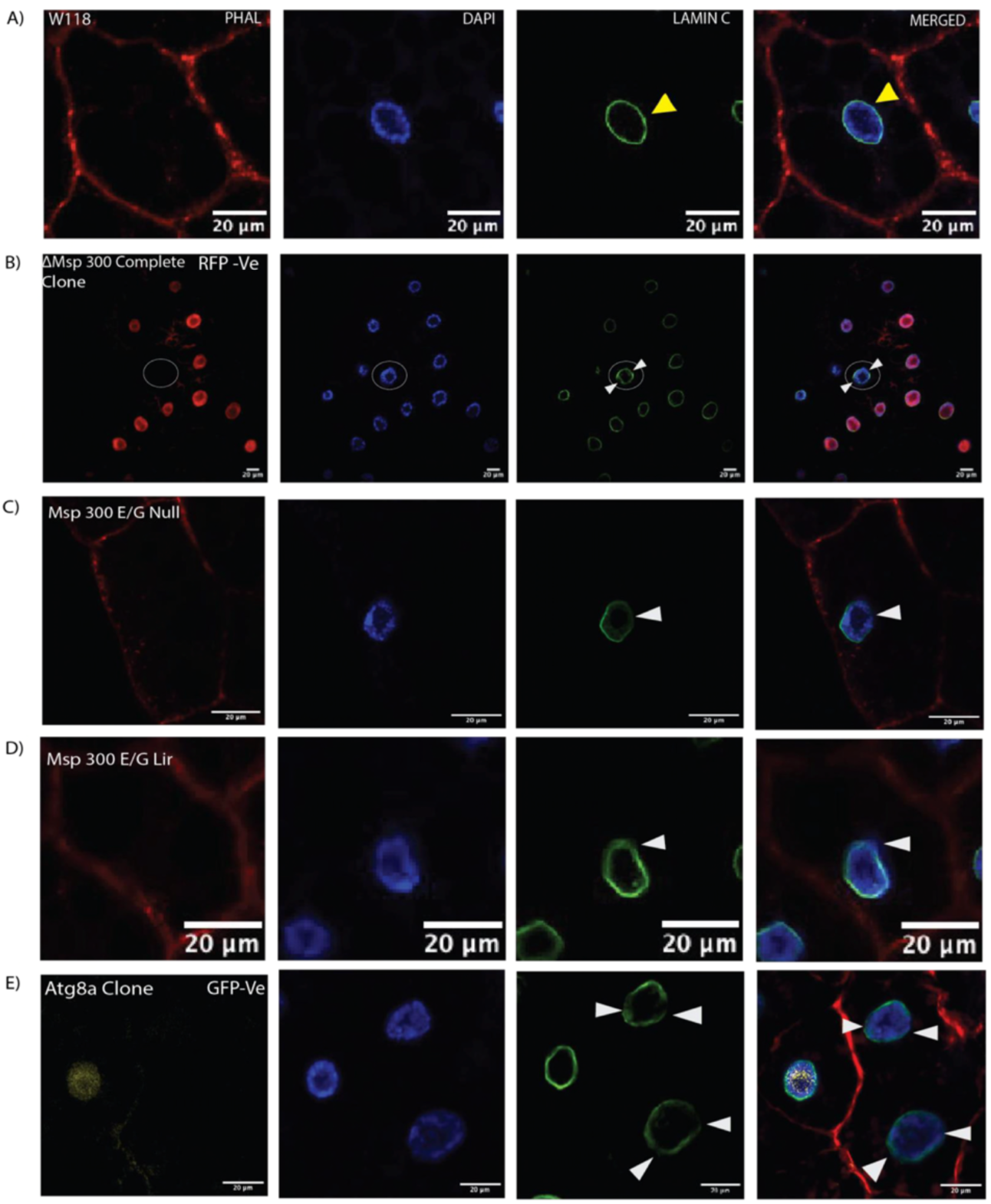
Disrupted Lamin C organization in Atg8a and Msp300 mutants. Representative images showing Lamin C organisation in fat body cells from late second-instar larvae. In wild-type cells, Lamin C forms a continuous ring surrounding the nucleus. In Msp300 null clonal cells (RFP-negative), Atg8a null clonal cells (GFP-negative) generated using the FLP-out system, Msp300 E/G null mutants, and Msp300 E/G LIR mutants, Lamin C displays an irregular distribution around the nucleus.

### Alpha-tubulin defects in Msp300 and Atg8a mutant

Nuclear Lamins help anchor and coordinate cytoskeletal networks; Lamin A/C deficiency or defects leads to an increased separation or altered positioning of the microtubule-organizing centre (MTOC) in cell (Lee *et al*., 2007). Microtubules act like railway tracks within the cell and span through the cytoplasm for transporting organelles, vesicles, and proteins to their proper destinations. The MTOC anchors these tracks (usually near the cell centre), allowing motor proteins (like dynein and kinesin) to haul the cargo (Vineethakumari *et al*., 2022). In our study, disruption of Lamins and misalignment of nucleus prompted us to look at alpha-tubulin localisation. In control line cells, we observed a neat ring of circumferential microtubules (MTs) around the nucleus with radial microtubules extending into the cell (**Figure. 18 A**). In Msp300 E/G Null mutant the circumferential MT ring was severely deformed and radial MTs were completely lost (**Figure. 18 B**). In Msp300 E/G LIR mutant circumferential MT ring was irregular while the radial MT network looked dense but irregular (**Figure. 18 C**). Most interestingly, in Atg8a mutant (KG07569) lacking functional copy of Atg8a, circumferential MT ring was completely lost, and radial MT network was completely haphazard (**Figure. 18 D**). Msp300 and Atg8a mutants did not have normal distribution of radial as well as circumferential MTs.

**Figure 18.**
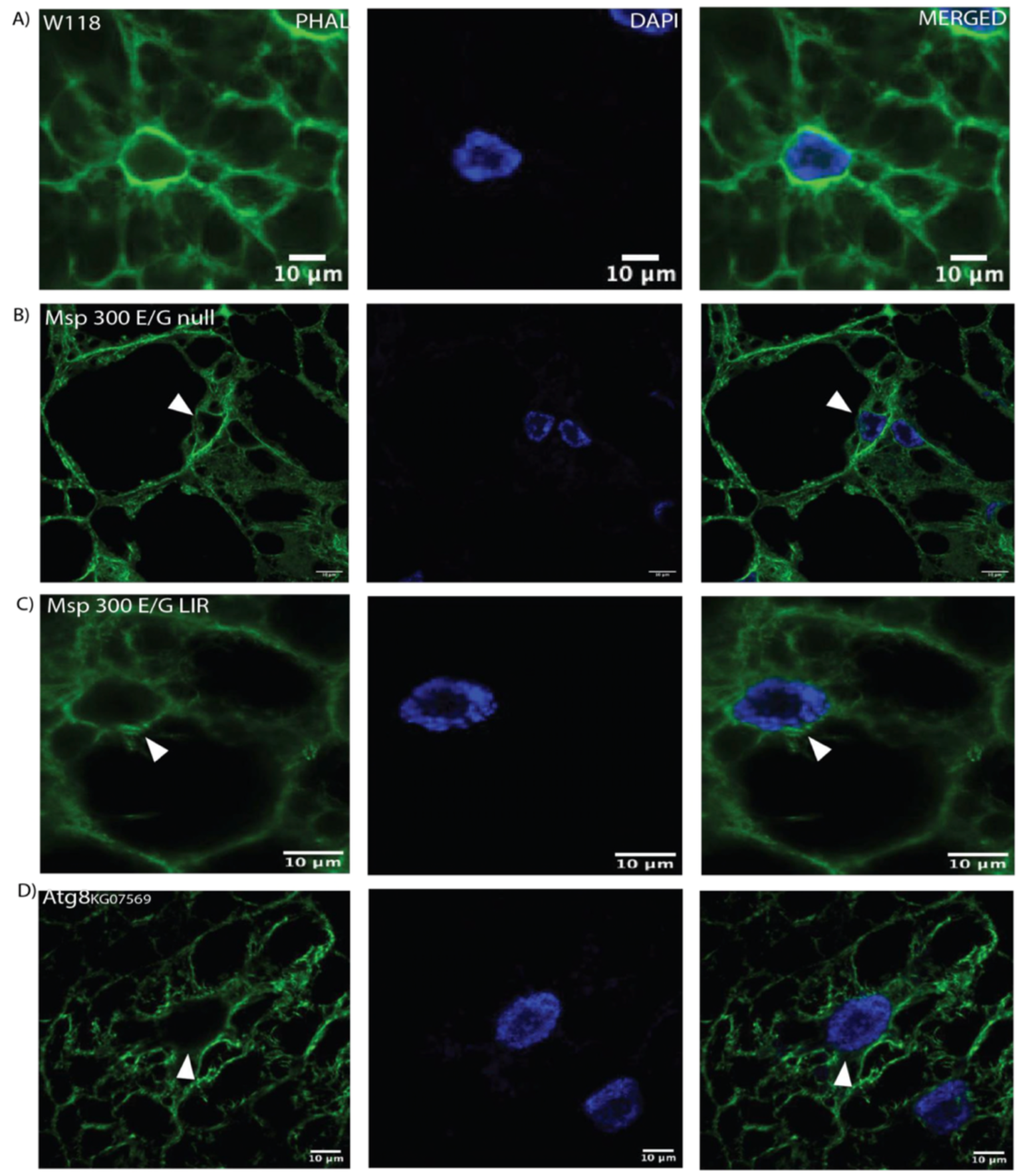
Alpha-tubulin defects in Msp300 and Atg8a mutant cells. Representative confocal image showing the normal distribution of microtubules (MTs) around the nucleus in w1118 fat body cells from instar larvae. Circumferential MTs form a ring around the nucleus, while radial MTs extend into the cytoplasm. **(B)** In Msp300 E/G null m circumferential MTs fail to form a ring (white arrow) and radial MTs are sparse or absent. **(C)** In Msp300 E/G LIR mutants, circumferent form an irregular ring that is impacted into the nucleus, while radial MTs are disorganised. **(D)** In Atg8a null mutants, both circumferentia MTs are in complete disarray. No distinct circumferential MT ring (white arrow) is observed, and radial MTs display an irregular pattern.

### Disrupted Endoplasmic Reticulum (ER) localisation and lipid metabolism in Msp300 and Atg8a mutant

Msp300 is a KASH domain protein that localises at ONM and all through the ER (van der Graaf *et al*., 2024). The interplay between tubulins and ER builds the ER structural framework within the cell. With the evidence of converging alpha-tubulin mutant phenotypes in Msp300 and Atg8a mutants, we wanted to check for ER localisation and function in Msp300 and Atg8a mutants. Calnexin is an integral membrane protein of ER that serves as molecular marker for ER (Paskevicius *et al*., 2023). Fat body is lipogenic tissue and lipid synthesis happens at the ER. We therefore, checked for ER functionality by looking at lipid synthesis in fat body of

In control line, like distribution of microtubules, calnexin has a nuclear ring corresponding to ER localised at ONM and the radial network corresponding to peripheral ER network extending throughout the cell. In Msp300 E/G null mutant ONM ring of calnexin was present however the radial network of ER was sparse In Msp300 E/G LIR mutant, both the nuclear network and the radial network were perturbed. In Atg8a mutant, the nuclear network of calnexin was present however the peripheral network of ER was sparse (**Figure. 19 A)**. Next, we performed western blot analysis to check for ER abundance in our mutants showing uncharacteristic ER distribution. Quite interestingly, control lines had less calnexin compared to all the other mutants (**Figure. 19 B).** In immunofluorescence lipid droplet (LD) analysis, we observed that control line had nucleus tightly packed with LDs around it. In Msp300 complete clonal mutant we observed RFP-ve cells had very few LDs, but they were bigger in size compared to control line. In Msp300 E/G null mutant we observed similar sparse LD phenotype, but LD were not any larger. In Msp300 E/G LIR mutant we observed slightly larger LDs tightly packed around in the nuclei. However, in the Atg8a mutant the LDs were prominently large (**Figure. 20 A**). Triacylglycerols (TAGs) are predominant form of lipids stored in fat bodies. We calorimetrically quantified the TAGs in early 3^rd^ instar larvae fed on normal diet. Quite surprisingly when quantified we observed less TAGs present in all the Msp300 and Atg8a mutant compared to the control line. In Atg8a mutant where we observed biggest lipid droplets, we found them to have least amount of TAGs. However, this difference within the three mutants was not statistically significant but collectively Atg8a and Msp300 mutants had defective lipid hemostasis (**Figure. 20 B)**.

**Figure 19.**
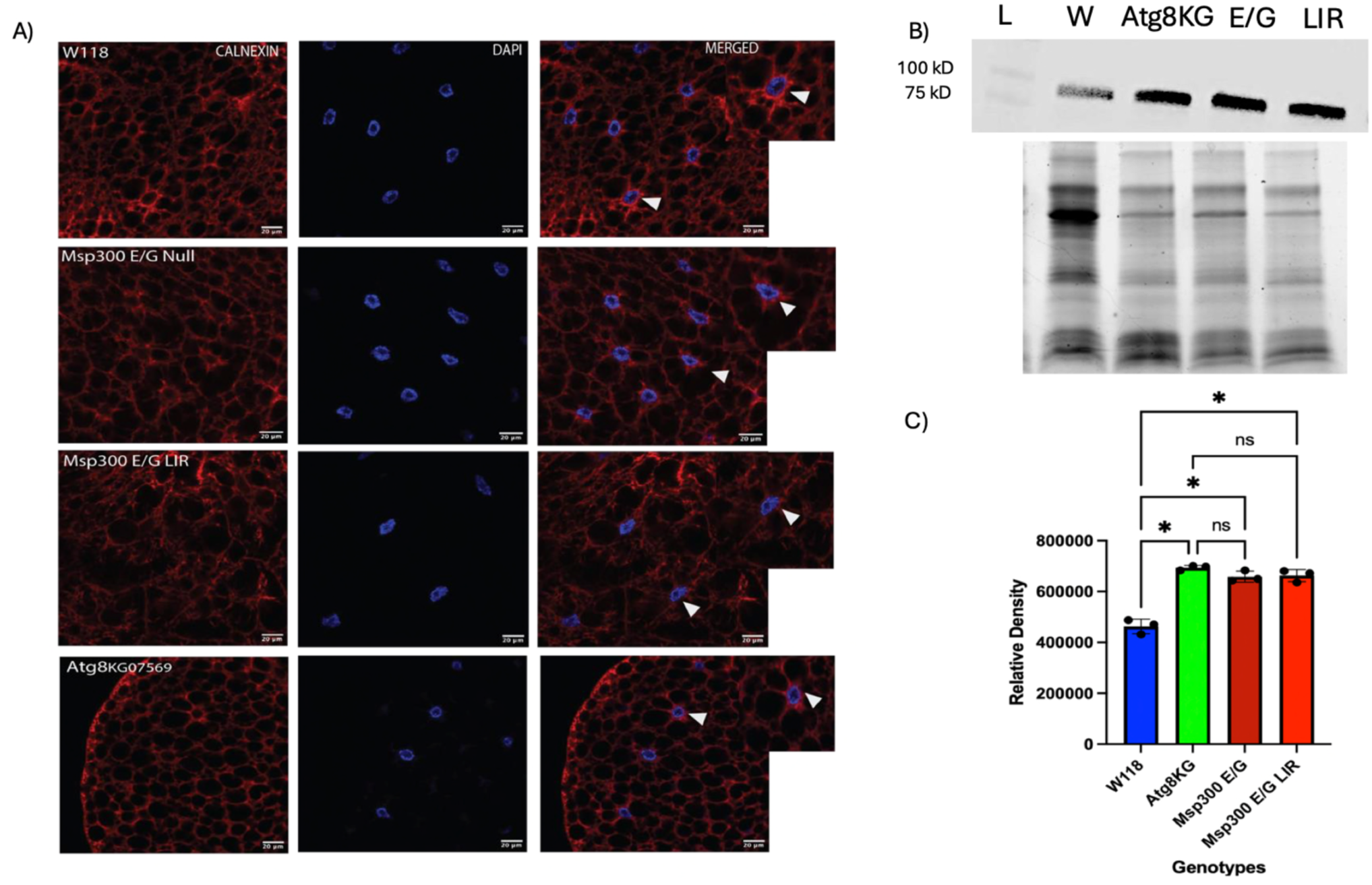
Disrupted ER organization in larval fat body cells of Msp300 and Atg8 mutants. **(A)** Representative images of the ER network in second-instar larval fat body cells with endoplasmic reticulum visualized in red (anti-calnexin antibody) and nucleus in blue (DAPI staining). In W1118, the ER network extends throughout the cell, with prominent signals at the cell periphery (Msp300-specific ER ring) and in the perinuclear region. The Msp300-specific ER ring is retained in Atg8-null mutants, although the ER network appears sparse. In Msp300 E/G null and Msp300 E/G LIR mutants, the ER ring is disrupted and distorted, with altered ER distribution. **(B)** Western blot analysis of calnexin accumulation in early third instar larvae of W1118, Msp300 E/G null, Msp300 E/G LIR, and Atg8 null mutants. Total protein staining was used for normalisation. (C) Quantification of calnexin levels. Data are presented as mean ± SD (n = 3). Statistical analysis was performed using one-way ANOVA followed by Šídák’s multiple-comparisons test. Significance is indicated as follows: ns (P > 0.05), * (P ≤ 0.05), ** (P ≤ 0.01), *** (P ≤ 0.001).

**Figure 20.**
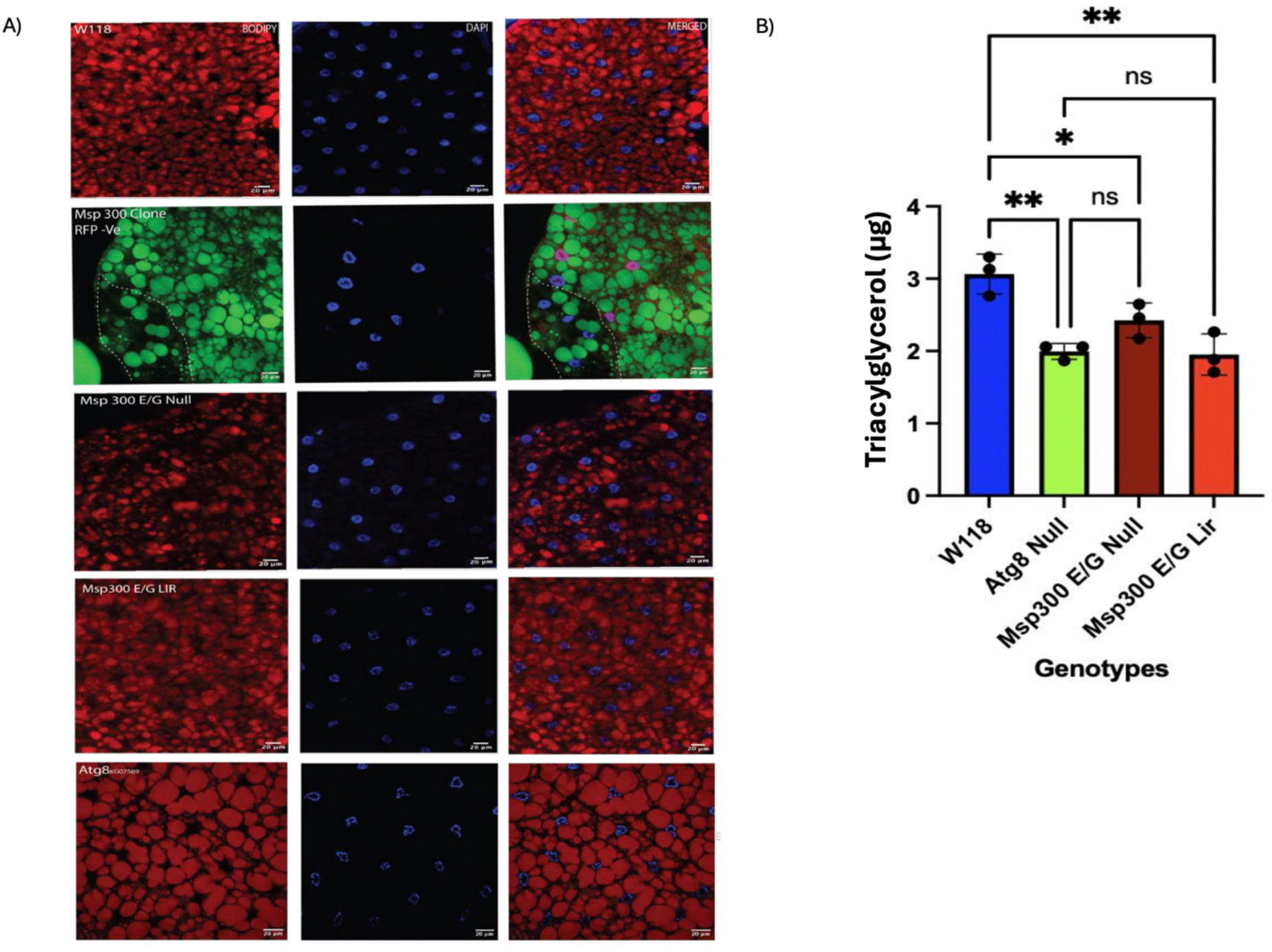
Defective lipid metabolism in Msp300 and Atg8a mutants. **(A)** Representative confocal images of fat body cells showing lipid droplets stained in red (BODIPY) and nuclei labelled in blue (DAPI) in w1118, Msp300 E/G null mutants, Msp300 E/G LIR mutants, Atg8a null mutants, and clonal Msp300 complete mutant cells (RFP-negative) using FLP-out system. Msp300 E/G null and Msp300 E/G LIR mutants exhibit irregular and sparse lipid droplets, whereas Atg8a null mutants display enlarged lipid droplets compared with w1118 fat body cells. All mutants indicate a defect in lipid homeostasis. **(B)** Triacylglycerol (TAG) colorimetric quantification shows significantly reduced TAG levels in Msp300 E/G null mutants, Msp300 E/G LIR mutants, and Atg8a null mutants. Data are presented as mean ± SD (n = 3). Statistical analysis was performed using one-way ANOVA followed by Šídák’s multiple-comparisons test. Significance is indicated as follows: ns (P > 0.05), * (P ≤ 0.05), ** (P ≤ 0.01), and *** (P ≤ 0.001).

### Autophagic defects in Msp300 mutants

With Atg8a being a central regulator of autophagy and Msp300 having only recently been shown to be required for autophagy and proteostasis in muscles (van der Graaf *et al*., 2024), it was imperative to investigate the involvement of Msp300 in autophagy and its interaction with Atg8a, with the aim of functionally characterising the nature of this interaction axis. Autophagy receptors such as Ref (2)P link ubiquitinated cargo to Atg8a on autophagosomes, ensuring their delivery to lysosomes for degradation. When autophagy is impaired, these substrates accumulate as ubiquitinated protein aggregates and so does the receptor that binds to such cargo, which is a hallmark observed in numerous human diseases. In this study, we checked for Ref (2)P and ubiquitin accumulation under basal conditions in early 3^rd^ instar larvae of Atg8a and Msp300 mutants. Atg8a mutants are autophagic mutants expected to have to high accumulation of both Ref (2)P and ubiquitinated cargo and thus acting as positive control for this part of the study. In immunofluorescence study, we found Msp300 E/G null mutant and Msp300 E/G LIR mutant accumulated high number of puncta/cell for both ubiquitin **(Figure. 21 A-A)** and Ref (2)P **(Figure. 22 A-A)** compared to the control lines. However, Ref (2)P and ubiquitin accumulation in both these mutants was less than Atg8a mutant. The western blot analysis of Ref (2)P corroborated the results from immunofluorescence and showed high Ref (2)P accumulation in both the Msp300 mutants (**Figure. 22 C-C**). The results pointed towards importance of Msp300 for basal autophagy.

**Figure 21.**
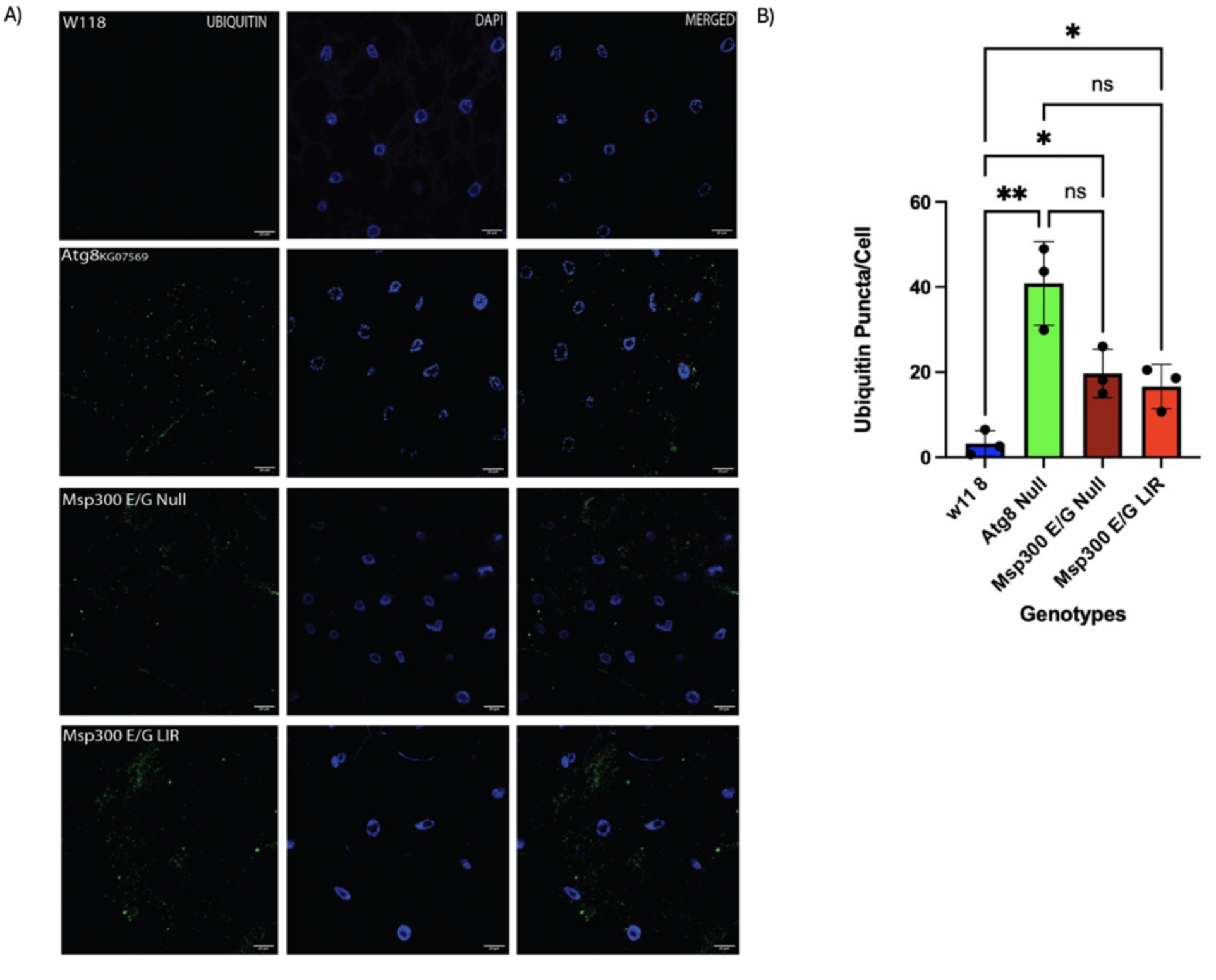
Mono and poly-ubiquitin accumulation in Msp300 and Atg8a mutants. **(A)** Representative confocal images of larval fat body cells showing F-actin (phalloidin, green), nuclei (DAPI, blue), and mono-and polyubiquitin (ENZ-ABS840, green) in W1118, Msp300 E/G null, Msp300 E/G LIR, and Atg8-null mutants. Increased ubiquitin accumulation is observed in Msp300 E/G null, Msp300 E/G LIR, and Atg8 null mutants compared with W1118, indicating impaired autophagic clearance of ubiquitin tagged cargo. **(B)** Quantification of ubiquitin accumulation. Data are presented as mean ± SD (n = 3). Statistical analysis was performed using one-way ANOVA followed by Šídák’s multiple-comparisons test. Significance is indicated as follows: ns (P > 0.05), * (P ≤ 0.05), ** (P ≤ 0.01), *** (P ≤ 0.001).

**Figure 22.**
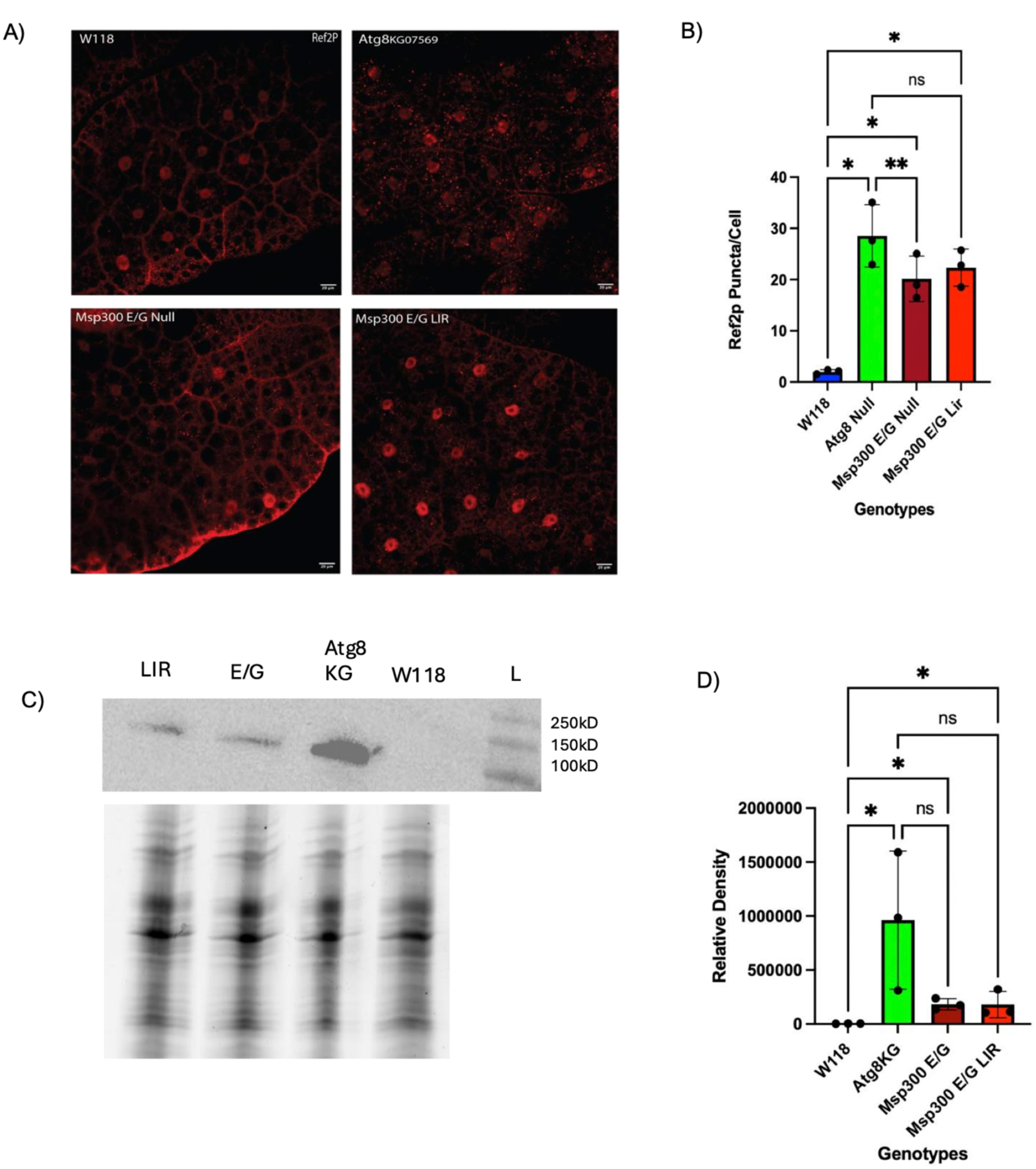
Ref(2P) accumulation in Msp300 and Atg8a mutants. **(A)** Representative confocal images of larval fat body cells for checking Ref(2P) accumulation in fat body cells of early third instar larvae in W1118, Msp300 E/G null, Msp300 E/G LIR, and Atg8-null mutants. Increased ubiquitin accumulation is observed in Msp300 E/G null, Msp300 E/G LIR, and Atg8 null mutants compared with W1118, indicating impaired autophagic clearance of ubiquitin tagged cargo. **(B**) Quantification of ubiquitin puncta/cell from immunofluorescence analysis. **(C)** Western blot for conforming Ref(2P) accumulation in late second instar larvae. Atg8 null used as a positive control for Ref(2P) accumulation and total lane protein normalisation for loading control. **(D)** Quantification of western blot. Data for are presented as mean ± SD (n = 3). Statistical analysis was performed using one-way ANOVA followed by Šídák’s multiple-comparisons test. Significance is indicated as follows: ns (P > 0.05), * (P ≤ 0.05), ** (P ≤ 0.01), *** (P ≤ 0.001).

In immunofluorescence study, we found Msp300 E/G null mutant and Msp300 E/G LIR mutant accumulated high number of puncta/cell for both ubiquitin **(Figure. 21 A-A)** and Ref (2)P **(Figure. 22 A-A)** compared to the control lines. However, Ref (2)P and ubiquitin accumulation in both these mutants was less than Atg8a mutant. The western blot analysis of Ref (2)P corroborated the results from immunofluorescence and showed high Ref (2)P accumulation in both the Msp300 mutants (**Figure. 22 C-C**). The results pointed towards importance of Msp300 for basal autophagy.

### Autophagosomes accumulate in Msp300 E/G RNAi mutant

We performed immunofluorescence analysis using Atg8a-mcherry-gfp tandem marker which can distinguish between Atg8a in autophagosomes from Atg8a in autolysosomes (Kimura *et al*., 2007). The m-Cherry tag attached to Atg8a tracks the location of Atg8 while the GFP tag tracks specifically the quenching of GFP in acidic compartment which is helpful in tweaking out Atg8a in autophagosomes and Atg8a in autolysosomes. Under normal conditions, autophagosomes should progress to become autolysosomes, and GFP quenching should be observed. Under stress however, if autophagy is intact, we expect to see more autophagosomes and more autolysosomes and in confocal there would be both red and yellow signals. However, if autophagy is impaired and autophagosomes don’t mature, we expect to see accumulation of autophagosomes which in immunofluorescence would mean lingering of GFP signal alongside m-Cherry and we will see accumulation of yellow Atg8a. In our experiment we compared RNAi control lines and Msp300 E/G RNAi lines under basal conditions and under amino acid starvation when autophagy is up regulated. Autophagosomes were counted per cell using analyse particle plugin per cell wherein each puncta was added as region of interest (ROI) in ImageJ/Fiji (version 2.14.0/1.54p) task manager. Per puncta per cell GFP intensity was calculated using the mean intensity build in plugin. To decide GFP cut off, average mean intensity was recorded for 10 GFP -ve punctae from obvious GFP -ve background.

In the RNAi control line under basal conditions (**Figure. 23 A**), we observed only a few red puncta, representing autophagosomes. Upon amino acid starvation, there was a considerable increase in the accumulation of red puncta, as expected. Some of these puncta co-localised with green puncta, resulting in a yellow signal, while others showed quenched GFP fluorescence and therefore exhibited only a red signal (**Figure. 23 B**). In the Msp300 E/G RNAi line under basal conditions a lot more autophagosomes accumulated compared to the to control line (**Figure. 23 C**). Under amino acid starvation, there was a considerable increase in the accumulation of red puncta and green puncta, as expected. Most of these puncta co-localised with green puncta, resulting in a yellow signal, very few red puncta showed quenched GFP fluorescence (**Figure. 23 D**). Quantification of yellow (GFP⁺/mCherry⁺) or red-only (GFP⁻/mCherry⁺) puncta in control and Msp300 E/G RNAi condition showed control RNAi had 68% yellow puncta

**Figure 23.**
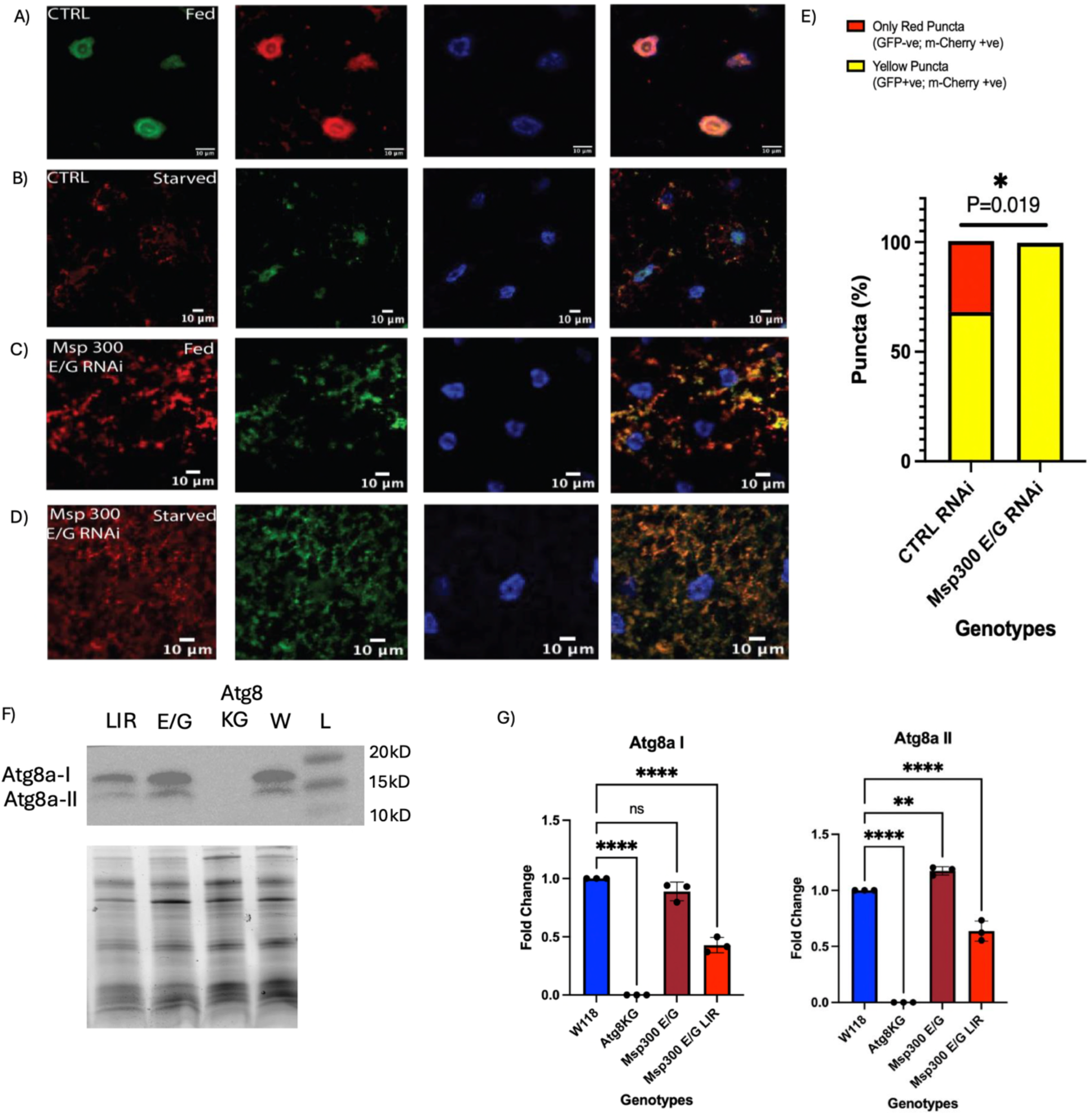
Impaired autophagosome degradation in Msp300 E/G RNAi mutants. **(A-D)** Autophagosome maturation in control and Msp300 E/G RNAi lines was assessed using the Atg8a-mCherry-GFP tandem reporter. Under fed conditions, control cells showed little Atg8a accumulation, whereas amino acid starvation increased Atg8a puncta, with GFP quenching indicative of autophagosome–lysosome fusion. Msp300 E/G RNAi cells showed increased Atg8a puncta under fed conditions, with a higher proportion retaining GFP signal. Following amino acid starvation, Atg8a puncta increased but showed limited GFP quenching, consistent with impaired autophagosome maturation. **(E)** Quantification of mCherry-positive puncta classified as yellow (GFP⁺/mCherry⁺) or red-only (GFP⁻/mCherry⁺) in control and Msp300 E/G RNAi under starved conditions. Data represent three independent experiments (n = 3). Statistical analysis was performed using two-way repeated-measures ANOVA followed by Šídák’s multiple-comparisons test. The genotype × puncta-type interaction was significant (P = 0.0119); adjusted pairwise comparisons were P = 0.0472 for yellow puncta and P = 0.0450 for red-only puncta. **(F)** Western blot analysis of Atg8a-I (unlipidated) and Atg8a-II (lipidated) in W1118, Msp300 E/G null, Msp300 E/G LIR, and Atg8a mutant. Atg8a mutant and W1118 were used as negative and positive controls, respectively. **(G)** Quantification of Atg8a levels from immunofluorescence and western blot analyses. Data are presented as mean ± SD (n = 3). Statistical analysis was performed using one-way ANOVA followed by Šídák’s multiple-comparisons test. Significance is indicated as follows: ns (P > 0.05), * (P ≤ 0.05), ** (P ≤ 0.01), *** (P ≤ 0.001).

(GFP⁺/mCherry⁺) and 14% red-only (GFP⁻/mCherry⁺) puncta (**Figure. 23 E**) while Msp300 E/G RNAi line had more than 99% yellow puncta (GFP⁺/mCherry⁺) (**Figure. 23 E**). We performed western blot analysis in Msp300 E/G null and Msp300 E/G LIR for checking autophagic flux by checking abundance of Atg8a-1(un-lipidated form) and Atg8a-II (lipidated Atg8a) (**Figure. 23 F**). In this experiment control line was used as positive control and Atg8a mutant was used as a negative control. We found that Msp300 E/G null mutant accumulated same amount of Atg8-I as positive control. while Msp300 E/G LIR mutant accumulated less Atg8-I. In case of Atg8-II, compared to the control line Msp300 E/G null mutant accumulated more lipidated Atg8a compared to control while the Msp300 E/G LIR mutants accumulated significantly Atg8-II. As expected, negative control Atg8a mutant did not have Atg8-I or Atg8-II (**Figure. 23 G**).

### Msp300 KASH containing isoforms co-localizes with Atg8a in cytoplasm and at the nuclear envelope

Msp300 typically comprises a series of spectrin repeats (SRs), a pair of calponin homology (CH) domains at the N-terminus, and a KASH domain at the C-terminus (14). However, isoforms E and G differ from this canonical structure: both lack the CH domains, while isoform G retains the KASH domain unlike isoform E which has another predicted transmembrane domain on N-terminus, and both contain spectrin repeats (Rey *et al*., 2021). The KASH domain, which includes a transmembrane helix and a short C-terminal region that anchors Msp300 to the outer nuclear membrane (Starr *et al*., 2005). Spectrin repeats are thought to form an extensive interaction platform with multiple binding sites, supporting the idea that SRs act as docking surfaces for signal transduction and integral membrane proteins (Liem *et al*., 2016). Autophagic defects observed in Msp300 mutants, combined with the presence of key domains in the isoforms of interest, prompt us to check if Msp300 and Atg8a interact at the nuclear periphery to facilitate autophagic activity around the outer nuclear membrane (ONM-phagy) during basal conditions and under stress. To investigate this mechanism, we crossed a GFP-tagged Msp300 reporter line (BDSC 59757) to an mCherry-Atg8a line driven by a fat-body-specific GAL4 driver. To induce a robust autophagic response, early 3^rd^ instar larvae were starved for 4 hours in a 20% sucrose solution (Scott *et al*., 2004).

Under fed conditions, Atg8a was predominantly nuclear with a few cytoplasmic puncta, and Msp300 is mostly localised to the outer-nuclear space as a nuclear ring with filamentous cytoplasmic network and very few cytoplasmic puncta (white arrows) which were potentially Msp300 associated with ER peripheral network (**Figure. 24 A**). Under amino acid starvation, we observed larger aggregates of Atg8a in cytoplasm and immediately around nucleus. Msp300 nuclear ring was visible and large punctae were present in cytoplasm and immediately around nucleus, that showed striking spatial overlap between Msp300 and Atg8a (**Figure. 24 B**). Under both the conditions i.e. fed and starved, we observed that this Msp300 ring was discontinuous and irregular in some nuclei (yellow rows) (**Figure. 24 A-A**). Colocalization analysis of Atg8a and Msp300 showed that, under starvation, a significantly larger fraction of Atg8a overlapped with Msp300 (Manders’ coefficient M1) and the Pearson’s correlation coefficient was higher than under fed conditions (**Figure. 24 C**).

**Figure 24.**
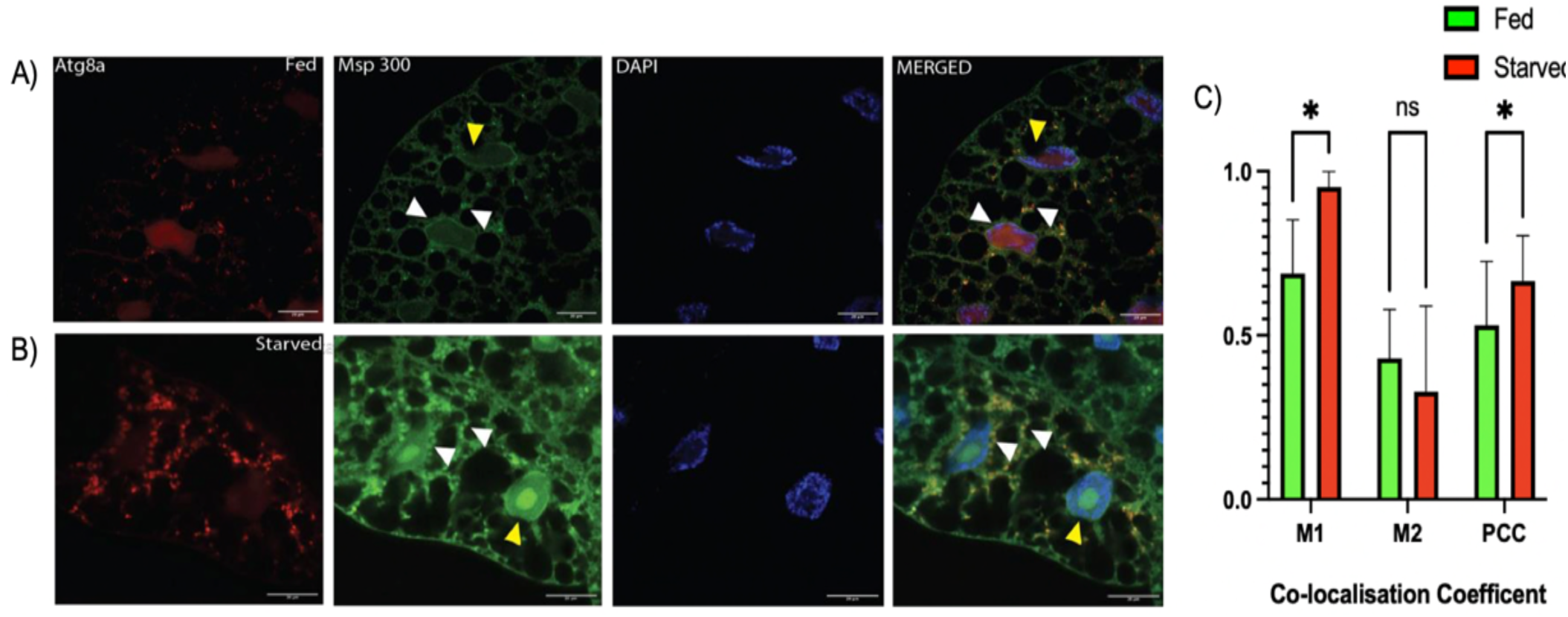
Co-localization of Msp300 and Atg8a at the nuclear periphery and in the cytoplasm under amino acid starvation. **(A)** Under fed conditions in the fat body of second-instar larvae, when autophagy is at basal levels, Atg8a is predominantly nuclear with some cytoplasmic puncta, and Msp300 is mostly localised to the perinuclear region as a perinuclear ring with a few cytoplasmic puncta (white arrows). **(B)** Under starvation-induced stress, large Atg8a aggregates are observed around the nucleus and within the cytoplasm while no nuclear Atg8a is detected. Msp300 is present in cytoplasm as large punctae and localised to the perinuclear region (white arrows). The Msp300 ring around the nucleus remains clearly visible, and cytoplasmic Atg8a aggregates show strong co-localisation with Msp300, indicating interaction between the two proteins when autophagy is upregulated under stress conditions. Under both fed and starved conditions, Msp300 perinuclear ring some nuclei show irregular Msp300 ring (yellow arrows). **(C)** Colocalization analysis of Atg8a and Msp300 under fed versus starved conditions. Thresholded Manders’ coefficients (M1, M2) and Pearson’s correlation coefficient (PCC) were calculated via Costes’ automatic thresholding (BIOP-JACoP, Fiji) and confirmed by Costes’ randomization (p = 1.00). Bars show mean ± SD; N = 23-40 images analyzed from 10 larvae/group. Unpaired t-tests with Welch’s correction, Holm-Šídák corrected for multiple comparisons. ns, P > 0.05; *P ≤ 0.05; **P ≤ 0.01; ***P ≤ 0.001.

### Dynamic remodelling of Msp300 outer nuclear membrane ring under stress

To further investigate the association between Msp300 and Atg8a during cellular stress, we fed second instar larvae with 24 um Tunicamycin and 100 mM etoposide for 24 hr, respectively.

Etoposide has been reported to cause DNA damage and invagination of nuclear envelope (Dellaire *et al*., 2009) while tunicamycin triggers unfolded protein response in endoplasmic reticulum (Chow *et al*., 2013) both of which trigger autophagy. Thereby we expected an increase in autophagic activity, marked by Atg8a accumulation and potential reorganization of Msp300 if it plays a role in alleviating stress. To test this, after dissecting drug fed larvae we carried out confocal microscopy on fixed fat body cells expressing m-cherry tagged Atg8a and Msp300 GFP (tags all KASH isoforms) and compared the signals with isoform-specific antibodies for Msp300 E and G to precisely determine the contributions of these isoforms.

In untreated control samples from KASH-GFP tagged 3^rd^ instar fat bodies, cytoplasmic Atg8 puncta were observed alongside a Msp300 nuclear ring. The cytoplasmic Msp300 puncta had spatial overlap with Atg8a puncta (**Figure. 25 A**). In DMSO-treated samples, the Msp300 nuclear ring was discontinuous, and Atg8a puncta strongly co-localised with cytoplasmic Msp300 puncta (**Figure. 25 A’**). Following etoposide treatment, H2Av was used as a marker of DNA double-strand breaks (**Figure. 25 B**). Atg8a and Msp300 puncta were observed, and the Msp300 nuclear ring was completely absent or discontinuous in cells. These Atg8a and Msp300 puncta completely spatially overlapped in the cytoplasm and around outer nuclear membrane (**Figure. 25 C**). Calnexin was used as a marker of the unfolded protein response following tunicamycin treatment (**Figure. 25 D**). Under these conditions, large cytoplasmic Atg8 puncta co-localised with Msp300 puncta within the cytoplasm and around the nucleus, and the Msp300 nuclear ring was completely lost or partially intact in fat body cells (**Figure. 25 E**).

**Figure 25.**
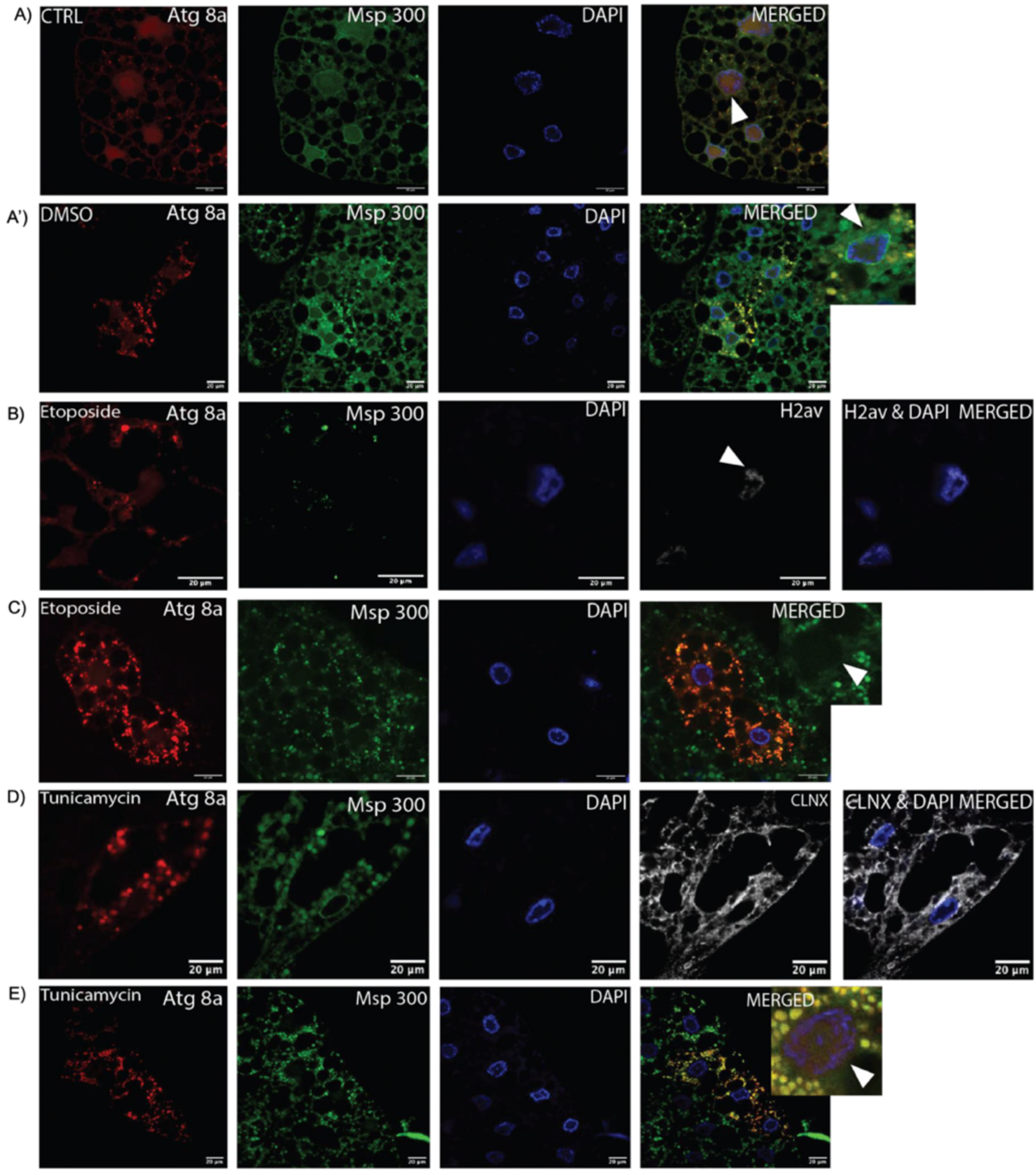
Stress-induced co-localisation of Msp300 and Atg8a at the nuclear periphery and in the cytoplasm, accompanied by loss of the Msp300 perinuclear ring. **(A-E)** Fat bodies of early third instar larvae using FLP-out system, fed with fly food containing, DMSO, 50 mM etoposide (DNA damage inducer) or 12 µM tunicamycin (ER stress inducer) for 24 h. **(A-A’)** Control without any drug and carrier. Etoposide and tunicamycin were prepared in DMSO; therefore, a DMSO control was used to monitor the effects of DMSO. Some cells had clear fully intact Msp300 nuclear ring in outer nuclear membrane while some cells had partly Mp300 ring (white arrow). **(B, D)** DNA damage from etoposide was confirmed using the double-strand break marker H2Av (white arrow), and ER stress from tunicamycin was confirmed using the Calnexin marker. **(C, E)** Both the treatments induced autophagy, as indicated by larger and more prominent Atg8a aggregates around the nucleus and within the cytoplasm and no one in the nucleus. Some fat body cells showed Atg8a-Msp300 co-localisation together with complete loss of the Msp300 nuclear ring (white arrows). These observations suggest stress-dependent remodeling of the outer nuclear membrane, potentially mediated by the Atg8a-Msp300 interaction through outer nuclear membrane-phagy (ONM-phagy).

We also wanted to study Msp300 E/G localisation and effect of stress on Atg8a and Msp300 E/G interaction. In DMSO-treated fat body explants, the Msp300 nuclear ring was present but discontinuous. Large cytoplasmic Atg8a puncta were observed, but they did not co-localise with cytoplasmic Msp300 E/G puncta, despite their proximity **(Figure. 26 A)**. Following etoposide treatment, large cytoplasmic Atg8a puncta were observed, whereas large Msp300 E/G puncta even though present did not co-localise with Atg8a puncta **(Figure. 26 B)**. In contrast, tunicamycin treatment resulted in large cytoplasmic Atg8a puncta that spatially overlapped with Msp300 E/G puncta, accompanied by complete loss of the Msp300 E/G nuclear ring and interestingly Msp300 E/G localised quite strongly to the nucleus. **(Figure. 26 C)**. This experiment specifically examined the E and G isoforms of Msp300. Interestingly; etoposide treatment did not induce the expected co-localisation of Atg8a puncta with cytoplasmic Msp300 E/G puncta in cytoplasm or outside the outer nuclear membrane. In contrast, tunicamycin treatment, which induces the unfolded protein response, resulted in strong co-localisation of Atg8a puncta with Msp300 E/G puncta in cytoplasm which is potentially Msp300 associated with peripheral ER network. These findings suggest that Msp300 KASH isoforms could be responsible for basal ONM and ER homeostasis while Msp300 E/G isoforms only associate with peripheral ER network and may be responsible for mediating stress-induced remodelling of ER network.

**Figure 26.**
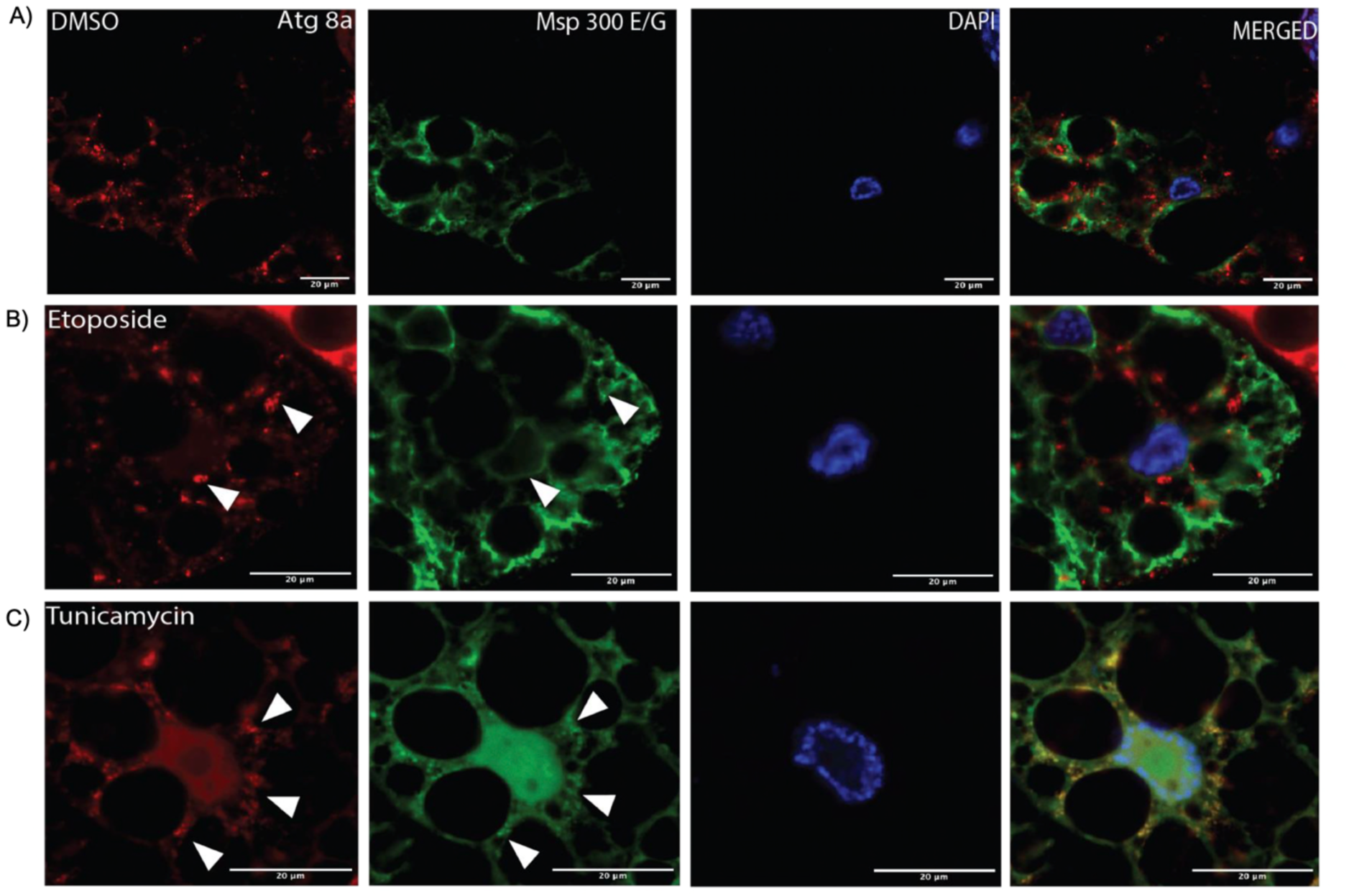
Stress-induced co-localisation Msp300 E/G isoforms with Atg8a and loss of the Msp300 nuclear ring. Immunohistochemistry was performed on flpout mCherry Atg8a expressing larvae treated with etoposide or tunicamycin ex-vivo and labelled using Msp300 E/G specific antibodies. (A) Etoposide and tunicamycin were prepared in DMSO; therefore, a DMSO control was used to monitor the effects of DMSO. The Msp300 nuclear ring was present but irregular. No Atg8a and Msp300 E/G co-localisation was observed at the nuclear periphery, although some cytoplasmic Msp300 E/G puncta spatially overlapped with Atg8a. (B) Following etoposide treatment, the Msp300 E/G nuclear ring was present but irregular. No Atg8a and Msp300 E/G co-localisation was observed at the nuclear periphery or between Atg8a and the cytoplasmic Msp300 E/G puncta (white arrows). (C) Following tunicamycin treatment, the Msp300 E/G ring was completely lost. Atg8a puncta strongly co-localised with cytoplasmic Msp300 E/G puncta (white arrows). The loss of the Msp300 E/G nuclear ring was consistent with stress-induced structural remodeling of the outer nuclear membrane. These findings suggest that stress triggers dynamic reorganisation of Msp300 E/G isoforms at the nuclear envelope, potentially mediated by Atg8a through ONM-phagy (outer nuclear membrane autophagy).

### Intact Msp300 nuclear ring but lost Atg8a interaction in Msp300 E/G LIR mutant

From the AlphaFold analysis and Y2H interaction study, we anticipated interaction between Atg8a and Msp300 E/G isoform to be mediated by the LIR domain and if LIR is mutated this interaction would potentially be lost. However, we wanted to check in adipocytes of early 3^rd^ instar larvae if Msp300 E/G still localise to ONM and ER after the LIR mutation. We were also interested to know if Msp300 E/G specific puncta and Atg8a co-localised i.e. are still in proximity to be still able to interact. We found that LIR mutation did not affect the localisation of Msp300 E/G isoform at ONM specific nuclear ring and in peripheral ER network **(Figure. 27 A)**. However, very interestingly under etoposide induced stress as observed before (**Figure. 26 B**) for Msp300 E/G isoform in control background no Msp300 E/G specific punctae were observed and Atg8a puncta even though present around ONM and ER network did not co-localise with Msp300 E/G isoform specific puncta (**Figure. 27 B**). Under tunicamycin stress, Msp300 E/G isoform specific puncta were observed which did not co-localise with Atg8a puncta (**Figure. 27 C**) unlike observed previously in control background (**Figure. 26 C**). These observations were in line with the results from AlphaFold analysis and Y2H analysis. ONM Msp300 E/G isoform specific nuclear ring was evidently irregular and very patchy under both the treatment conditions.

**Figure 27.**
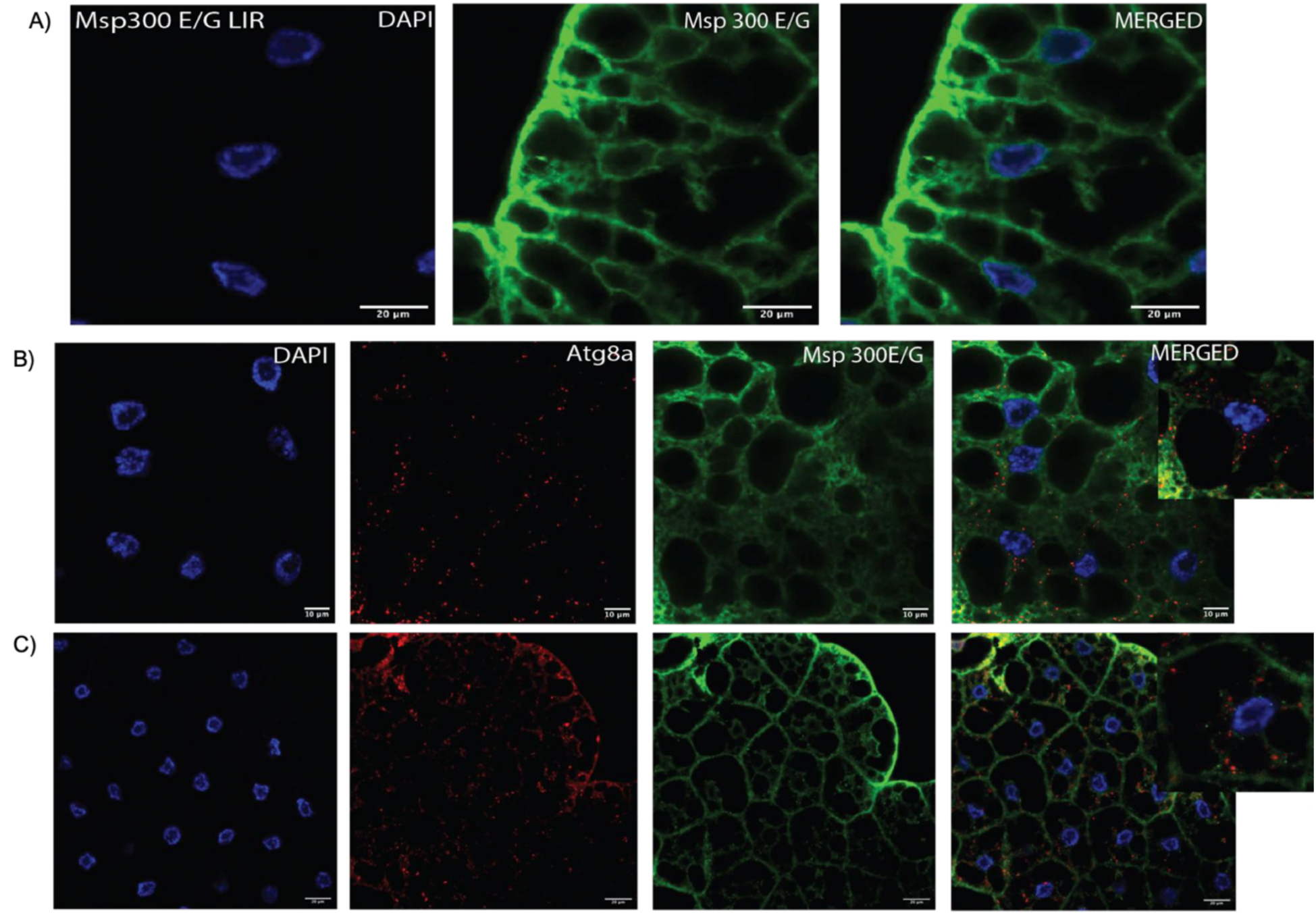
Intact Msp300 nuclear ring but lost Atg8a interaction in Msp300 E/G LIR mutant. Immunohistochemistry was performed on Msp300 E/G LIR mutant 3^rd^ instar larvae treated with etoposide or tunicamycin ex-vivo and labelled using Atg8a and Msp300 E/G specific antibodies. **(A)** Msp300 E/G specific LIR mutation doesn’t affect localization of Msp300 E/G isoform to ONM and ER peripheral network. **(B)** Following etoposide treatment, the Msp300 E/G ONM nuclear ring was present but irregular at times. No Msp300 E/G puncta were seen. No Atg8a and Msp300 E/G pucnta co-localisation was observed. **(C)** Following tunicamycin treatment, ONM nuclear ring remodeling was evident. Msp300 E/G pucnta were seen within the cell which did not co-localise with Atg8a puncta.

## Discussion

This study found a novel AlphaFold 3 predicted and Y2H validated interaction between Atg8a, a core autophagy regulator and LINC complex protein-Msp300 isoform E and G. This work for the first time shows a novel role of Atg8a in modulating nuclear position in fat body cells of 3^rd^ instar *Drosophila* larvae. We looked at cytoskeleton and nucleoskeleton defects in Msp300 and Atg8a mutant to understand how Atg8a affects nuclear positioning. There was no possible way to study the CH and KASH less Msp300 isoform E mutants in isolation from KASH containing Msp300 G, due to the common exon usage. Therefore, the mutants created in this study are referred to as Msp300 E/G Null mutants and Msp300 E/G LIR mutants. Very interestingly, we observed converging phenotypes of disrupted Lamin Dm0, Lamin C and alpha tubulin in our Atg8a and Msp300 E/G mutants which explained nuclear positioning defects in both Msp300 and Atg8a mutant. In these Msp300 and Atg8a mutant, we also evidenced, ER mis localisation and ER accumulation and disrupted ER synthesised TAGs. Such defects have not been reported before in Atg8a and they strongly suggest co-operation between Msp300 and Atg8a in maintaining nuclear morphodynamics.

Larval adipocytes are in state of miotic arrest under normal steady state conditions (Guarner *et al*., 2018) and Msp300 E/G have been shown to independently organise ncMTOC in *Drosophila* 3^rd^ instar larvae (Zheng *et al*., 2020). Microtubules are anchored perinuclearly by Msp300 at their minus ends while their plus ends stabilised at cell periphery by actin cytoskeleton and spectraplakin bridge (Sun *et al*., 2019; Zheng *et al*., 2020). These microtubules have been reported to move pre-fusion autophagosomes toward the nucleus/ncMTOC to locate lysosomes for degradation (Jahreiss *et al*., 2008). Another study has shown significant reduction in the number of mature autophagosomes when microtubules are impaired, but this has been shown to not affect their life span or their fusion with lysosomes (Fass *et al*., 2006). A recent study on Msp300 has reported autophagic defects in Msp300 complete mutant. Thereby in this study, we wanted to explore if Msp300 E/G isoforms coordinate with Atg8a for autophagy. We observed ubiquitin and Ref2(P) accumulation in Msp300E/G Null mutants and Msp300 E/G LIR mutants under steady state conditions. Msp300 E/G RNAi mutant additionally showed impaired autophagosome maturation. This observation clearly consolidated the requirement of Msp300 E/G for maintaining basal autophagy.

Msp300 is localised at ONM and in peripheral ER network and has been recently shown to not only co-localise with the LAMP1 lysosomal marker but also in confocal light scanning microscopy (CLSM), appeared to be present within the autolysosome (Kucińska *et al*., 2023). In our study, we observed Msp300 isoform E and G puncta associated with Atg8a only under ER stress while interestingly no colocalisation of punctae were observed under control conditions or under DNA damage stress even though the filamentous network of Msp300 E/G and Atg8a spatially overlapped. Unlike the KASH containing isoforms of Msp300 which we found to strongly co-localise with Atg8a under basal autophagy, amino acid starvation, DNA damage and ER stress. We also encountered a unique remodelling of Msp300 specific nuclear ring in steady state conditions as well as under stress. The evidence for this study suggests that Msp300 specific nuclear ring that tethers nucleosketon and cytoskeleton is constant under quality control surveillance. Atg8a has been shown to support oligomerization of ER-Phagy SARs (Mochida *et al*., 2020). In our study, we observed loss of Msp300 isoform E and G puncta in LIR mutants which could be due to lost interaction between Atg8a and Msp300 isoform E/G which in turn caused loss in receptor oligomerisation. Atg8a interacts with SARs at several membrane and organellar contact sites. Given that LC3A has previously been shown to stabilise microtubules (Faller *et al*., 2009) and GABARAP also associates with microtubules (Wang *et al*., 2000), it is plausible that Atg8a at the ONM and ER, contributes to microtubule stabilisation and supports the formation and maintenance of the cellular cytoskeletal framework. However, it must be noted that their seems to be a bidirectional relationship between autophagy and microtubules. On one hand pre-autophagosome are transported via microtubules, when the intracellular clearance pathways are impaired, damaged organelles and protein aggregates accumulate, increasing ROS levels and disrupting microtubule networks. This cellular stress can ultimately compromise autophagic function, further exacerbating the decline in cellular homeostasis. Our study strongly suggests that the LINC complex protein Msp300 (in an isoform specific manner) recruits Atg8a, a core autophagy component, to maintain nuclear and ER homeostasis. We propose that under steady-state conditions, the outer nuclear membrane (ONM) and ER-resident protein Msp300 acts as an architectural tether that interacts with Atg8a to stabilise the microtubule network and maintain membrane contacts sites (MCS) required for ER organisation. This Atg8a under basal or induced autophagy becomes lipidated and accumulates at the selective phagophore assembly site (PAS), where it promotes the stable sequestration and tight docking of bulky cargo by facilitating the oligomerisation of selective autophagy receptors.

## Conclusion

In conclusion, Atg8a and Msp300 isoform specific interaction drives ER and nuclear homeostasis in adipocytes of *Drosophila melanogaster*. LINC complex protein Msp300 recruits’ autophagy protein Atg8a at ONM and ER through LIR domain mediated interaction. This interaction helps in maintaining membrane contacts sites which form stable microtubules cytoskeleton. In abrogated LIR mutants where the Atg8a and Msp300 interaction is lost this results in impacted nucleoskeleton and cytoskeleton, which in turn lead to ER homeostasis defects and autophagic defects observed in this study. This work also shows very interesting phenomenon of ONM specific Msp300 ring remodelling under steady state and under stress suggesting a novel role of Msp300 in maintaining cellular homeostasis.

## Acknowledgement

The authors gratefully acknowledge Hybrigenics Services (Paris, France) for their technical assistance with yeast two-hybrid (Y2H) screening and WellGenetics Inc. (Taipei, Taiwan) for their expert assistance in generating the CRISPR-edited Msp300 E/G null and Msp300 E/G LIR mutant fly lines. We sincerely thank Gábor Juhász (University of Budapest), Péter Lőrincz (Eötvös Loránd University) and Timothy Megraw (University of Florida) for their generous provision of the transgenic fly lines used in this study. We also extend our sincere thanks to Timothy Megraw for providing the Msp300 E/G isoform-specific antibody used in this work. This work was supported by the Biotechnology and Biological Research Council (BBSRC) and University of Warwick funded Midlands Integrative Biosciences Training Partnership (MIBTP).

## Author Contribution

Shristi Sharma conducted the research, generated, analysed, visualised the data, and drafted this manuscript for thesis. Ioannis P. Nezis along with Shristi Sharma conceptualised the study. Ioannis P. Nezis provided supervisory oversight and project administration.

## General Discussion

In this study, we investigated the interaction between Atg8a, a core autophagy protein, and Msp300 E/G, a LINC complex protein, to determine its role in autophagy and nuclear morphodynamics. Selective autophagy receptors and adapters proteins that interact with Atg8a contain short linear motifs-LIR motifs (LC3-interacting region motifs), LC3 recognition sequences (LRS), or Atg8-interacting motifs (AIM), which are required for recognition and interaction of SARs with Atg8-family of proteins (Pankiv *et al*., 2007; Ichimura *et al*., 2008; Noda *et al*., 2010, Rehman *et al*., 2022). LIR motif-containing proteins bind via their LIR motif to the hydrophobic pocket 1 (HP1) and hydrophobic pocket 2 (HP2) of the LIR docking site (LDS) of the respective Atg8 protein (Johansen and Lamark, 2020; Rehman *et al*., 2022). Amino acid K48A and Y49A of Atg8a are imperative for LIR-LDS interaction. Point mutations (K48A and Y49A) within the hydrophobic pockets of LDS of Atg8a have been shown to abolish interactions with LIR dependent SARs (Johansen and Lamark, 2020; Rehman *et al*., 2022). In this study, we used AlphaFold 3 analysis for prediction of interaction between Msp300 isoform E and isoform G, respectively with Atg8a. For both the isoforms, AlphaFold predicted high confidence interaction between Atg8a within the amino acid residue 46-55 and Msp300 E/G within17-residue stretch (residues 8967-8983). The residues of interaction were particularly interesting because for Atg8a, LDS falls within the range of amino acid that showed interaction (46-55aa) and for Msp300 E/G iLIR software predicted a 6aa LIR motif (residues 8966-8971) within the sequence predicted to interact by AlphaFold. We also used Y2H analysis to confirm a strong interaction between Msp300 E/G and Atg8a proteins. Mutation analysis confirmed this interaction is facilitated by a 6aa LC3-interacting region (LIR) motif. This LIR motif is a common feature of a many selective autophagy receptors that bind Atg8a. In this study, we point mutated 2 aa (EDYVII → EDAVIA) that are important for formation of hydrophobic pockets where Atg8a docks to the receptor for recruiting autophagy machinery. This LIR mediated Y2H interaction with high PBS (predicted biological score) between Atg8a and Msp300 isoform E/G laid foundation for the present study and provided first crucial evidence linking the autophagy machinery to nuclear-cytoskeletal machinery of cell.

Msp300 is reportedly known to regulate nuclear spacing, attachment of astral microtubules and positioning of ER and mitochondria in *Drosophila* (Elhanany-Tamir *et al*., 2012; Morton *et al*., 2025). It has 11 spatiotemporally regulated isoforms of which, isoforms E and G have been shown to cooperate for anchoring myonuclei in larval skeletal muscle (Rey *et al*., 2021) and in organising the ncMTOC and nuclear positioning in the fat body of *Drosophila* larvae (Morton *et al*., 2025). In our study, mutation of the LIR motif in Msp300 produced a similar phenotype of loss of nuclear positioning in the adipocytes of 3^rd^ instar *Drosophila* larvae. Interestingly, loss of Atg8a also produced a phenotype resembling that of the LIR mutant and the Msp300 E/G null mutant. Quantitative analyses of clonal fat body cells confirmed significant differences in both nuclear centrality and circularity in cells deficient in Atg8a compared with control type cells. For this experiment we used Atg8a RNAi line (109654) from Bloomington. In a recent study it was found that chromosomes carrying TRiP insertions into attP40 can confer dominant nuclear clustering effects due to closeness of attp40 site to Msp300. Due to which in certain genetic backgrounds, nucleus clustering effects are observed in RNAi lines but not due to true RNAi instead because of Msp300 insertional mutation (Van der Graaf *et al*., 2022). However, for Atg8a RNAi, VDRC KK line used in this study, RNAi this is uniquely integrated into cytogenetic position 30B which does not share proximity to attP40 (25C) docking site.

LINC component tethers nucleoskeleton and cytoskeleton. In our study, we observed nucleus localisation defects, the nature of which cannot be understood without looking at the key components of nucleoskeleton and cytoskeleton. In our Atg8a mutants and Msp300 (E/G Null and E/G LIR) mutants we distinctly observed Lamin Dm0 and Lamin C defects. Such defect in Lamin localisation and reduction of Lamins around nucleus upon Msp300 knockdown has been reported previously (Shaw *et al*., 2022). Lamin C predominantly interacts with heterochromatin and plays an important role in gene regulation (Gurudatta *et al*., 2010). In contrast, Lamin Dm0 contributes to the attachment of nuclei to the microtubule organising centre (MTOC), facilitating nuclear migration through its cooperation with microtubules and Klarsicht (Patterson *et al*., 2004). We therefore checked microtubule arrangement in our Atg8a and Msp300 E/G Null and E/G LIR mutants.

Msp300 isoform E/G have been shown to organise alpha-tubulin independent nucleation of ncMTOC in fat body cells of *Drosophila* which is responsible for nucleus positioning (Zheng *et al*., 2020). In this study, Msp300 isoform E was shown to be sufficient for ncMTOC assembly independently of Msp300 isoform G. In our study we observed microtubules organisation defect in our Atg8a and Msp300 (E/G Null and E/G LIR) mutants which corroborated the nuclear positioning defects. In context of autophagy microtubules facilitate cellular trafficking including autophagosomes, maintains organelle integrity, forms molecular scaffolds, and establishing diffusion barriers on membranes (Cao *et al*., 2026). Cellular organelles such as ER depend on microtubular network for spatial distribution. ER tubules rely on microtubule network for transport bidirectionally (Wang Zheng *et al*., 2023). In our present work we found disorganised calnexin, accumulation of calnexin in western blot and unusual TAG produced on ER in our Atg8a and Msp300 (E/G Null and E/G LIR) mutants. We also observed autophagy defects in Atg8a and Msp300 (E/G Null and E/G LIR) mutants in our study. Ref (2)P, a receptor of autophagy and ubiquitin, a tag for degradation ready cargo, accumulated under basal conditions. Atg8a driven autophagy machinery utilizes microtubules tracks for transport of autophagic cargo to designated compartments. If the tracks are not continuous defects in autophagy is expected.

This can impact cellular ability to maintain ER homeostasis not only by not being able to clear damaged ER but also not being able to distribute ER tubules along the microtubular network which can manifest as disorganised ER in immunohistochemistry and accumulation of ER resident protein in western blot. Our western blot experiment showed high level of lipidated Atg8a in Msp300 E/G null mutants which is outcome of high autophagic flux but considered together with our rests from fluorescence microscopy of tandem Atg8a marker which showed accumulation of yellow Atg8a puncta upon induction of amino acid starvation, shows impaired clearance of autophagosomes. One previous study has linked Msp300 to autophagy and protein turnover within larval body wall muscles. This study suggests that ER-localised Msp300 plays a role in maintaining the cytoskeletal contacts required for autophagosome completion (van der Graaf *et al*., 2024). However, the underlying mechanism and the way in which Msp300 interacts with the autophagic machinery remain unclear. Atg8a has been shown to support oligomerization of ER-Phagy SARs (Mochida *et al*., 2020). In our study, we observed loss of Msp300 isoform E and G puncta in LIR mutants which could be due to lost interaction between Atg8a and Msp300 isoform E/G which in turn caused loss in receptor oligomerisation. Atg8a interacts with SARs at several membrane and organellar contact sites. Given that LC3A has previously been shown to stabilise microtubules (Faller *et al*., 2009) and GABARAP also associates with microtubules (Wang *et al*., 2000), it is plausible that Atg8a at the ONM and ER, contributes to microtubule stabilisation and supports the formation and maintenance of the cellular cytoskeletal framework.

In our Msp300 E/G LIR mutants however, low levels of Atg8a-I and Atg8a-II were observed in western blot. This was very interesting taken together with Ref(2)P and ubiquitin accumulation points towards a more systemic issue of basal level autophagy and therefore importance of LIR motif and Atg8a-Msp300 interaction for basal autophagy. Msp300 E/G LIR mutants show more drastic defects perhaps because Msp300 E/G isoforms are structurally present and localise at their usual location but are unable attend cellular duties because of lack of functional LIR motif. Failure in interaction could perhaps result is low cytoplasmic pool of Atg8a. However, in Msp300 E/G Null mutant both the isoforms are absent and under such conditions there is perhaps functional redundancy from the other isoforms contributing to relatively more cytoplasmic Atg8a. Our Atg8a tandem marker indicated clear autophagosome accumulation in Msp300 E/G RNAi under fed as well starved conditions which is indicative of general autophagy defect in these mutants.

To further develop understanding of role of Msp300 in autophagy we checked localisation of both Atg8a and Msp300 within the fat body. We found evidence of Msp300 co-localising with Atg8a within the cytoplasm under basal conditions. Interestingly, this co-localization is enhanced under amino acid starvation stress. We hypothesised Msp300interacts with Atg8a for modulating outer nuclear membrane and ER homeostasis. Upon Etoposide treatment, we observed large puncta of Atg8a in cytoplasm and around ONM which co-localised with Msp300 KASH isoform but not with E/G isoform. However, upon tunicamycin treatment, we observed the puncta of Atg8a which co-localised with both Msp300 KASH isoforms and Msp300 E/G isoform.

However, in LIR mutant we did not observe any co-localisation between Atg8a and Msp300 E/G isoform which supported the evidence from AlphaFold and Y2H interaction studies. In all the cases ONM Msp300 ring was found to be remodelled under stress with the ring appearing either partially or completely lost. It was very interesting to see the ONM-ring of Msp300 remodelling under steady state conditions. ER-stress and unfolded protein response have been shown to cause disassembly of the LINC complexes via asymmetric autophagy of the nuclear envelope involving selective autophagy of Nesprin, mammalian orthologue of Msp300 (Kucińska *et al*., 2024). From the present study it became evident under ER stress, only Msp300 E/G localised with Atg8a while under DNA damage stress and in steady state Msp300 KASH isoforms localised with Atg8a.

The convergent mutant phenotypes observed in Atg8a and Msp300 (E/G null and E/G LIR) mutants, with the loss of either Msp300 isoforms or Atg8a suggest a functional relationship between these proteins. This relationship when disrupted, the mutants displayed faulty cytoskeletal and nucleoskeletal components potentially due to their improper turnover from autophagic defects. They also displayed disrupted ER homeostasis evidenced by calnexin localisation defects, its accumulation in mutant and TAG levels. Autophagy has been shown to stabilise microtubules by degrading SCG10, a microtubule-destabilising protein, and to promote axon regeneration in neurons (He *et al*., 2016). In our study we found high confidence interaction of Msp300 E/G with Atg8a in the fly brain with current evidence it is possible Msp300 utilises Atg8a for stabilising microtubules in the neurons. In a recent bioRxiv preprint in ageing neuron defective autophagy is suggested to cause cytoskeletal collapse due to high reactive oxygen species (ROS) from accumulating damaged and dysfunctional mitochondria (Gupta *et al*., 2026). However, in this study we didn’t probe this further in the brain of the fly. LC3A mammalian orthologue of Atg8a has been shown to stabilise microtubule networks by decreasing microtubule dynamicity (Faller *et al*., 2009). GABARAP, another mammalian orthologue of Atg8a closely related with about 73% to 75% sequence identity, based on its structural conformation and head to tail oligomerisation, has been predicted to promote tubulin polymerization and facilitate GABA_A_ receptor clustering (Coyle *et al*., 2002). GABARAP has also been shown to associate with the cytoskeleton both in vitro and in vivo and suggested to play a role in the interaction between GABA_A_ receptors and the cytoskeleton (Wang *et al*., 2000). Their seems to be a bidirectional relationship between autophagy and microtubules. Proteostasis and organellar quality control is a coordinated action between repair and clearance of damaged proteins and dysfunctional organelles. Impairments in the intracellular clearance mechanisms result in the progressive accumulation of damaged organelles and aggregates of aberrant proteins which cause disruption of microtubules tracks from all the accumulated waste and high ROS, to then causing autophagic defects. In similar fashion mitochondrial dysfunction has been shown to drive disassembly of microtubules which in turn stop mitochondrial turnover and autophagosome clearance (Arduíno *et al*., 2013). On the other hand, alterations in the microtubule network themselves disrupt autophagosome clearance. Microtubules drive the transport and localization of autophagosomes by relying on “dynamic instability” and the “tubulin code” (Cao *et al*., 2026). Therefore, it is evident that their seems to be a mutual reliance of one process on the other and a healthy alliance between autophagy and microtubules is important for regulating cellular homeostasis. Our study strongly suggests that the LINC complex protein Msp300 recruits Atg8a, a core autophagy component, to maintain nuclear and ER dynamics by maintaining the necessary membrane and cytoskeletal contact sites between cellular organelles and potentially even stabilising the microtubules.

### Limitations of the study and Future Perspective

This study provides the first report of novel interaction between Atg8a and nesprin homologue, Msp300. The interaction and its LIR motif dependency was predicted by AlphaFold analysis with high confidence. This was study further backed by Y2H mutation analysis with high predicted biology score in head and 3^rd^ instar larvae. The confocal microscopy demonstrated colocalisation of the two proteins in perinuclear and cytoplasmic region. Together, these findings provide complementary structural, experimental and cellular evidence supporting a potential interaction between the two proteins. Nevertheless, the results from AlphaFold and Y2H are predictions while results from colocalisation only point at spatial proximity and do not establish direct binding, and further validation may be required to confirm the Atg8a-Msp300 interaction and of specific residues involved in this. The study shows the role of Atg8a in nuclear positing by potentially keeping the microtubule tracks clear and degrading ROS produced from damaged cellular organelles which destabilise microtubules. However, this involvement of Atg8a in nuclear placement could be explored further and especially the disorganised lamins. Since lamins form scaffolds for chromatin it would be worth checking chromatin compaction not only in Atg8a but also in Msp300 mutants.

Atg8a interacts with SARs at several membrane and organellar contact sites. GMAP recruits Atg8a at the Golgi, while Msp300 potentially recruits Atg8a at the outer nuclear membrane (ONM) and endoplasmic reticulum (ER). Given that LC3A has previously been shown to stabilise microtubules, it is possible that Atg8a performs a similar function at these sites, contributing to microtubule stabilisation and thereby supporting the formation and maintenance of the cellular cytoskeletal framework. Therefore, it would be interesting to study interaction between Atg8a, microtubules and microtubules associated proteins for exploring this potential role of Atg8a.

In this study, we also observed defects in ER localisation and function under normal fed conditions (i.e. under basal autophagy), as assessed by calnexin localisation and TAG localisation and quantification, respectively, in Atg8a and Msp300 mutants. This study pointed toward role of this interaction in ER homeostasis, quality control and importantly, ER-Phagy. The Immunofluorescence assay showed visually larger LDs in Atg8a and Msp300 mutants.

However, upon TAG estimation we observed less TAG in these flies. This is a classic sign of accumulation of toxic cholesteryl esters in fat bodies this study would benefit from quantifying cholesteryl esters alongside TAG for CE:TAG ratio in the fat bodies. The CE/TAG ratio determines whether a lipid droplet remains fluid or forms liquid-crystalline (LC) phases and high CE/TAG ratios are linked to disease.

Since Msp300 is also an ER resident protein. In this context, it would be important to tweak apart ER-phagy and general autophagy related defects in our Msp300 mutants. Atg8a mutants on the other side are expected to be defective in different kinds of selective autophagy because SARs-Atg8a is imperative for most selective autophagy pathways because it ensures efficient cargo-membrane tethering, phagophore elongation, and proper vesicle closure. To definitively determine whether both the Msp300 mutants have disrupted ER-phagy, general autophagy, or both, it will be important to look at the physical “stalling” of the autophagosome maturation or autolysosome formation using tandem GFP-mCherry reporters. For ER-phagy, the GFP-mCherry-Rtnl1(Reticulon 1) or GFP-mCherry-Atl (Atlastin) reporter can be used as these ER resident proteins which are also SARs have been extensively studied for their role in ER-phagy. Because these sensors change from yellow (autophagosomes) to red (autolysosomes), any block in the pathway will cause a buildup of yellow puncta. If Msp300 mutants are ER-phagy impaired, the GFP-mCherry-Rtnl1 or GFP-mCherry-Atl line will show a massive collection of yellow puncta. If it is a general autophagy mutant, GFP-mCherry-Atg8a will stall as yellow instead. If it is an impaired for both the Rtln/Atl and Atg8a tandem reporter lines will both show the accumulation of yellow puncta. It is interesting to check if Msp300 mutant is strictly an ER-phagy mutant, knocking down a known ER-phagy player in that background would not have an additive effect of yellow puncta accumulation. Similarly, it will also be important to knock down a core autophagy machinery gene, such as Atg1 to check its effect on the total autophagosome load. Studying Atg8a marker alongside Msp300 tandem tag and looking at their co-localisation with LAMP1. Msp300 tandem puncta co-localised with Atg8a and LAMP1 puncta would be strong evidence for the role of Msp300 E/G in ONM and ER-phagy. Perinuclear space is known to be increased under ER stress, perinuclear space measurement in our Msp300 and Atg8a mutants using electron or super resolution microscopy would provide an additional layer of evidence for Msp300 mutants are ONM-phagy deficient. In our work, we observed LIR mutants were unable to co-localise with Atg8a, but we still observed Msp300 E/G specific ONM-ring remodelling which does not align with the idea of LIR motif mediated Atg8a and Msp300 interaction responsible for ONM remodelling. However, ER remodelling is evident, and these interactions would be helpful in also tweaking apart Atg8a-Msp300 mediated ER and ONM remodelling.

To further consolidate, Msp300 isoform E/G is an autophagy receptor, it would be interesting to perform a protease protection assay. In this assay protease treatment would destroy all the cytoplasmic protein but proteases cannot cross the lipid bilayer. If Msp300 isoform E/G are autophagy receptor and is engulfed within the autophagosome it wouldn’t be digested and these samples when run on western blot would be positive for Msp300 isoform E/G.

In our autophagy maturation study using Atg8a tandem marker, we observed accumulation of autophagosomes in Msp300 E/G RNAi mutant under normal as well as starved conditions.

However, treatment with bafilomycin A1 treatment is imperative for making a concrete conclusion about impaired autophagic flux. If there is no additive effect of bafilomycin A1on autophagosome accumulation and in lipidated Atg8a-II levels that would be a clear indication of impaired flux. In these mutants, it would also be important to study why we observe autophagosome accumulation. This could be achieved by checking for autophagosome fusion or acidification defect using LysoTracker Red. In the case of fusion defect, co-localisation study with Atg8a and LysoTracker Red will show plenty of bright LysoTracker-positive puncta that do not co-localise with Atg8a. This would prove that lysosomes are perfectly acidic and healthy; they just aren’t fusing with the autophagosomes. However, if the LysoTracker signal will disappear or dim drastically across the entire cell. This will prove that the lysosomes themselves have lost their proton gradient. In conclusion, this study is advancing the field of autophagy and the mechanisms of selective autophagy by proposing a new selective autophagy receptor that helps in maintaining cellular homeostasis under basal and under stress conditions.

## Notes

### Competing Interest Statement

The authors have declared no competing interest.

